# Drivers of metabolic density-dependence: how resource availability and conspecific cues affect phytoplankton metabolism

**DOI:** 10.1101/2025.10.26.684709

**Authors:** Ricardo Estevens, Anna Lena Heinrichs, Giulia Ghedini

## Abstract

Metabolism is density-dependent from unicellular to multicellular organisms. Understanding what drives metabolic suppression is important to explain population growth given the link between metabolism and biomass production. In the simplest scenario, metabolic suppression is caused by a reduction in resource availability with increasing population density. But both theory and experiments suggest that organisms can actively downregulate metabolism in crowded conditions. We experimentally disentangle the importance of resource competition and conspecific cues that signal crowding on the metabolism of three phytoplankton species of varying cell sizes at different growth phases. All species downregulated some aspects of their metabolism in response to cues; this response varied in strength but could not be explained by differences in species size. The addition of nutrients weakened and, in some cases, completely removed metabolic suppression, indicating that resource availability mediates responses to cues. Overall, respiration rates were more responsive to cues than photosynthesis, showing a differential regulation of processes of energy intake and expenditure depending on both resource availability and conspecific cues. These factors also led to rapid plastic changes in cell size possibly related to cell division and growth. Altogether, changes in size and metabolism indicate that cues can trigger self-regulatory adjustments that might limit growth, but these effects are modulated by nutrient availability and species traits not related to size. These results suggests that growth predictions solely based on resource availability might overestimate the rates at which organisms and populations grow, with important implications for how we describe species and community dynamics.

## Introduction

Body size and temperature are two main variables that affect metabolism but population density is another important factor. Both unicellular and multicellular organisms have lower metabolic rates in denser populations (DeLong et al. 2014). The generality of density-dependent metabolic rates suggests that these metabolic adjustments are important for fitness and population regulation (DeLong and Hanson 2009; Nadler et al. 2016; Malerba et al. 2017; Lovass et al. 2020). Clarifying the drivers of metabolic suppression has implications for many fundamental and applied questions related to species abundances and growth. Indeed, the density-dependence of metabolism often tracks that of production, both in populations and communities (Sibly et al. 2005; Hatton et al. 2015; 2024; Fant and Ghedini 2024).

What drives density-dependence? An obvious and likely driver is resource competition. Increases in population density strengthen (intraspecific) competition and reduce resource availability. Thus, lower metabolic and growth rates in denser populations can be a direct consequence of diminished resource intake (DeLong and Hanson 2009). However, metabolic suppression occurs even without a decrease in resource availability (i.e. when respiration rates are measured in the absence of resources) (DeLong et al. 2014; Lovass et al. 2020). This result suggests that there might be other sources of regulation beyond food availability.

One hypothesis is that density-dependence might partly stem from behavioural adjustments, such as when organisms actively respond to intraspecific signals (Abrams 2022). These signals might often take the form of chemical cues that are widespread across the tree of life (Wyatt 2014). Cues (infochemicals or by-products of metabolism) can mediate diverse interactions, such as mating, aggregation, pathogenicity, or prey–predator relationships in both plants and animals (Gjoni et al. 2020; Lagos and Herberstein 2017; Kuhlisch et al. 2024; Saha and Fink 2022). Therefore it is likely that organisms adjust their metabolic activity in response to chemical cues if these contain information about the surrounding environment. Previous work indeed showed that chemical cues from conspecifics influence metabolic rates, suppressing metabolism even in the absence of resources (Nadler et al. 2016; Lovass et al. 2020). But the relative importance of resource competition and chemical communication for metabolic regulation is unclear.

Why would organisms reduce their metabolism in response to conspecific signals if resources are not limiting? This response seems counter-intuitive: higher metabolic rates increase competitive ability and growth (Auer, Salin, Anderson, et al. 2015; Pettersen et al. 2020), so we would expect organisms to perform at their maximum when resources are not limited.

However, mechanisms of self-regulation beyond direct resource competition are often needed in theoretical models to explain the diversity of natural systems (Barabás et al. 2017; Gavina et al. 2018; Hatton et al. 2024). It is possible that organisms, from microbes to multi-cellular eukaryotes, show some social behaviour or response to group living that regulates growth before reaching food limitation (Flux 2001; Miller et al. 2013; Ross-Gillespie and Kümmerli 2014). These metabolic adjustments might anticipate reductions in resource availability, more than responding to it, and might avoid population crashes – so they might have some fitness benefit. This type of metabolic flexibility (e.g., switching from high to low metabolic rates) might thus maximise fitness across environments (Auer, Salin, Rudolf, et al. 2015; Ghedini and Marshall 2023). Nonetheless, empirical tests of how organisms adjust their metabolism in response to population crowding are rare, since it is difficult to disentangle the effect of social cues from those of food limitation.

We attempt to disentangle these factors using marine phytoplankton as a model system. Chemical communication is common in marine organisms, including phytoplankton (Pohnert et al. 2007; Kuhlisch et al. 2024). Chemical cues mediate phytoplankton sexual reproduction (Moeys et al. 2016) and programmed cell death (Gallo et al. 2017), but little is known about their effects on phytoplankton metabolism. Our goal is not to identify a specific molecule but rather to quantify how cues from conspecifics present in spent media (e.g., metabolites and by-products of metabolism released by organisms in their environment) affect metabolic rates and determine if their effects are mediated by nutrient availability. We test the generality of this idea on three species differing in cell size (*Dunaliella tertiolecta*, *Tetraselmis sp.* and *Nannochloropsis granulata*) to understand if metabolic suppression is also related to this morphological trait, given its correlation with nutrient uptake and storage (Hillebrand et al. 2022). We exposed populations to media that differed in the presence of nutrients and cues, using an orthogonal design, at different timepoints during their growth curve (Figure 1) since the production of cues might be mediated by population growth stage (e.g. crowding signals might be more abundant when cells approach stationary phase) (Lazazzera 2000). After allowing cells to acclimate, we measured three aspects of metabolism (photosynthesis, post-illumination respiration, and dark respiration) as a function of population biovolume; and cell size to determine if changes in morphology accompany metabolic responses. We hypothesise:

*H1. Metabolic density-dependence:* all three species will have density-dependent metabolic rates (a relationship < 1 between population metabolic rates and biovolume on a log-log scale) because nutrients become limiting as populations grow. However, this density-dependence will be relieved when populations are resuspended in media with newly added nutrients (i.e. slopes ∼ 1 since cells are of very similar sizes within a population).
*H2. Cues effects:* for populations of equivalent biovolume and growth stage, cues from conspecifics will suppress metabolic rates but this response will be alleviated by the addition of nutrients (changes in intercepts; Figure 1c).
*H3. Cell size effects:* responses to both nutrients and cues will be stronger in smaller species because they have little capacity for nutrient storage and, thus, might be more responsive to nutrient pulses (a greater surface area to volume ratio should increase nutrient uptake).

**Figure 1.**
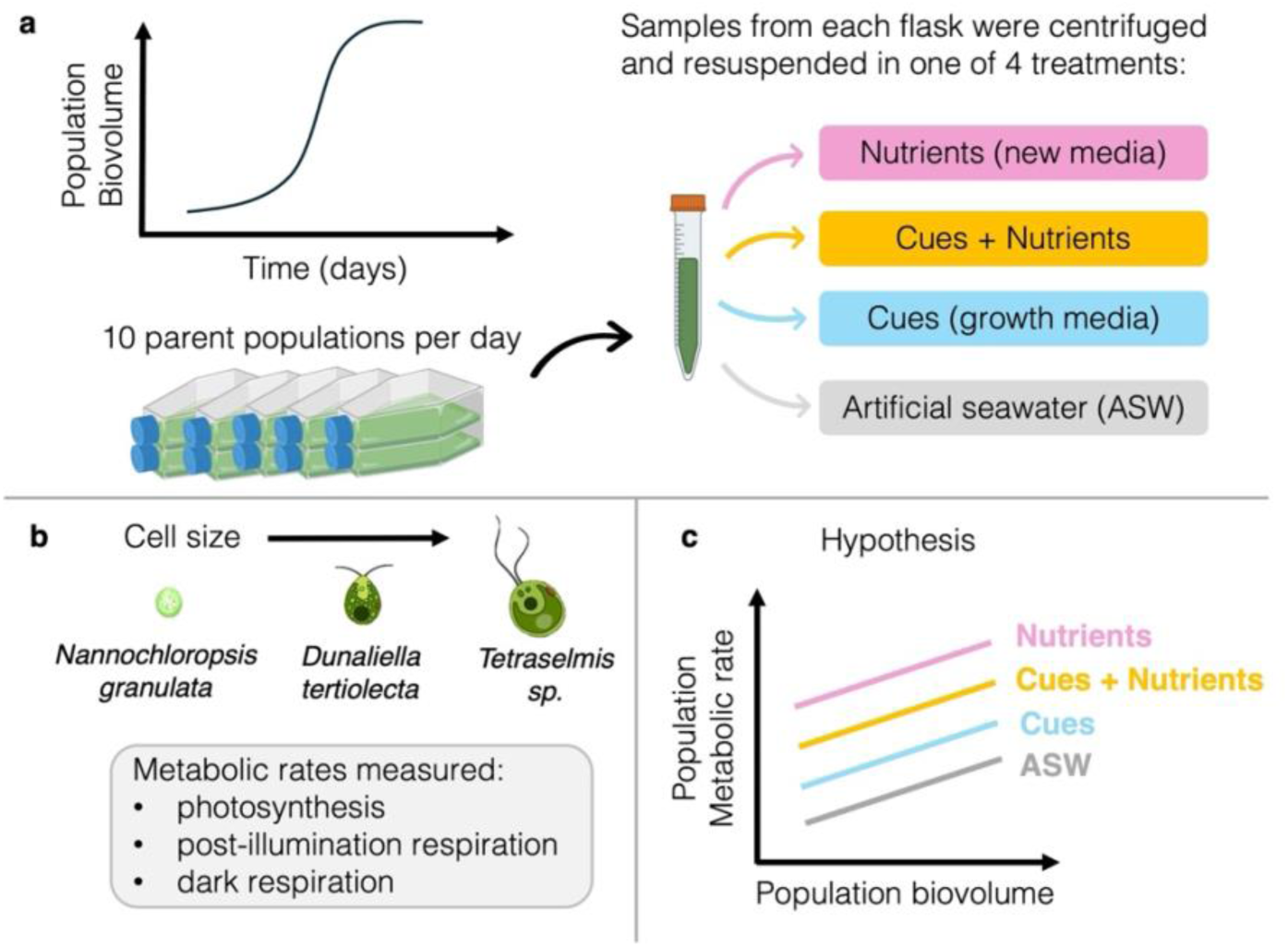
Experimental design. (a) To disentangle the effects of nutrient availability and conspecific cues on metabolic suppression, we measured the metabolic rates of three phytoplankton species as they increased in biovolume over time. Each day, we sampled ten flasks (parent populations). The content of each flask was divided into four sub-samples, centrifuged to obtain a pellet and then resuspended in one of four media treatments manipulating nutrients and cues in an orthogonal design (see also Box 1 for treatment details). After 3 hours (and also 30 minutes for *Dunaliella*), we measured population metabolic rates and cell size. (b) We repeated these measurements on three species that differed in size. (c) We hypothesize that, across the entire biovolume range, populations exposed to nutrients alone should have the highest rates, while those exposed to artificial seawater should have the lowest rates. We further hypothesize that cues reduce metabolism, but the addition of nutrients weakens their effect. Created in BioRender.

## Materials and Methods

To quantify the relative role of nutrient availability and conspecific cues on metabolic suppression, we grew three species of marine phytoplankton in monocultures and measured their photosynthesis and respiration rates after resuspending the cells in media that differed in the presence of nutrients and cues (Box 1). We repeated these measurements on different growth days, from exponential to stationary phase, to investigate if the response of the species to nutrients and/or cues depends on their growth phase. We used the species *Nannochloropsis granulata* (RCC438), *Dunaliella tertiolecta* (RCC6), and *Tetraselmis sp.* (RCC126) acquired from the Roscoff Culture Collection, France. We chose these species because they respond well to centrifugation, which was essential for this experiment, and are of different sizes, which we hypothesise might mediate their response to nutrients and cues.

### Box 1.

**Media Treatments**

1) “**nutrients**”: cells were resuspended in an equivalent volume (10 mL) of new artificial seawater media (the same formulation as the growth media) that thus contained nutrients but none of the cues present in the spent media;
2) “**cues + nutrients**”: we kept the cells in their original media to maintain the conspecific cues (the sample was shaken to resuspend the cells) but we added nutrients in an equivalent concentration to the growth media;
3) “**cues**”: cells were resuspended in the same spent media where they were growing; this media contained the cues released by the cells and the remaining nutrients that were not yet consumed (this amount of nutrients decreases over time as populations approach stationary phase);
4) “**artificial seawater (ASW)**”: cells were resuspended in an equivalent volume (10 mL) of artificial seawater that had no nutrients or cues.

Before starting the experiment, each species was cultured in artificial seawater F media (ASW F) (Guillard and Ryther 1962) in 1L Schott bottles with 15 minutes of filtered atmospheric air bubbling every hour, light intensity set at 50 µmol photons m^-2^s^-1^ and a temperature of 20°C. The same artificial seawater media was used throughout the experiment to avoid any confounding from variability in seawater nutrient concentration. Artificial seawater was prepared by mixing 35 g of salt (Instant Ocean, Aquatic Systems) into 1L of distilled water before sterilization. For both the initial cultures and the experiments detailed below, the photoperiod was set at 12h/12h light/dark cycle for *Dunaliella* and 14h/10h for the other two species as they were tested at different seasons.

### Experimental set-up

For each species, we cultured 120 parent populations with an initial biovolume of 5 × 10^4^ µm^3^/µl in T75 flasks (Sarstedt, Germany) with a total volume of 50 mL per flask. Each sampling day, we used ten parent populations while the remaining were left untouched. From each of these ten, we created four new populations (in 10 mL falcon tubes) which were then exposed to the media treatments, resulting in 10 replicates per treatment. To do this, each falcon tube was centrifuged at 3500 rpm (5000 rpm for *Nannochloropsis* due to its smaller size) for 10 minutes to precipitate the algal cells (AFI-C200RE, France). The algal pellets from each falcon tube were then resuspended in one of four types of media (Figure 1, Box 1).

Cells were left in these conditions for 3 hours before measuring population metabolic rates, biovolume and cell morphology. To determine if metabolic responses to the media treatments changed across growth phases, we sampled each species on different growth days, measuring ten new parent populations each day (12 sampling days for *Dunaliella* and *Nannochloropsis*; 8 for *Tetraselmis* because samples became contaminated after day 12).

### Cell morphology, density, and total biovolume

Before measuring metabolism, we collected 1 mL from each falcon tube which we fixed with 1% Lugol’s iodine (Carl Roth, Germany). Ten µl of this sample were loaded onto a Neubauer counting chamber (Marienfield, Germany) to take 20 images with light microscopy at 400x magnification (Olympus IX73 inverted microscope). All images were analysed using ImageJ and Fiji software (version 2.0) which recorded the length and width of each cell. We used these measurements to calculate the average cell volume (µm^3^ based on prolate spheroid shape for *Dunaliella* and *Tetraselmis* and a spherical shape for *Nannochloropsis* (Hillebrand et al. 1999), cell density (cells/µl) and biovolume (µm^3^/µl; multiplying cell density and average cell size) for each falcon tube.

### Metabolic rates

After 3 hours of exposure to the media treatments, we measured changes in oxygen levels in 5 mL vials filled with the content of each falcon tube using a respirometry system (PreSens Sensor Dish Reader, Precision Sensing, Germany) previously calibrated with 0% and 100% oxygenated artificial seawater. To prevent carbon limitation during the measurement, we added 50 µl of sodium bicarbonate to a final concentration of 2 mM. The samples were measured for 20-25 minutes under the light (at the same intensity used to grow the cultures, 50 µmol photons m^-2^ s^-1^) to quantify photosynthesis, followed by 40-45 minutes in the dark to measure respiration rates. We discarded the first three minutes of light and dark to allow samples to acclimate to the respective conditions. The first 15 minutes of the dark phase were used to calculate post-illumination respiration rates which showed a faster decline in oxygen levels; the remaining ∼20 minutes of dark were used to calculate dark respiration rates.

Since samples were not axenic, we corrected metabolic rates for bacterial respiration (i.e. changes in oxygen levels measured in blanks). The blanks were filled with the supernatant obtained by centrifuging the samples at 5000 rpm for 10 minutes to precipitate the algal cells. Each sampling day we had two sets of seven blanks: “fresh media blanks” were those collected from (and used to correct) samples re-suspended into new media (“nutrients” and “ASW” which should have very few if no bacteria at all), and “control blanks” were those collected from (and used to correct) samples re-suspended in their growth media which had more bacteria (“cues + nutrients”, “cues”).

The photosynthesis and respiration rates (μmol O_2_/min) of each sample were calculated as:

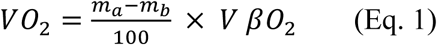

where m_a_ is the rate of O_2_ change in oxygen levels of the sample (min^-1^), m_b_ is the mean O_2_ level of the respective blanks (min^-1^), V is the sample volume (0.005 L) and βO_2_ is the O_2_ capacity of air-saturated seawater at 20℃ and 35 ppt salinity (225 µmol O_2_/L).

### Additional experiment on Dunaliella tertiolecta

We include an additional assessment which we performed on *Dunaliella tertiolecta* before the main experiments. In this initial assessment we acclimated samples for 30 minutes instead of 3 hours and did not include the treatment “cues + nutrients” (Box 1). We report these results because they offer insights on the plasticity of metabolic rates and whether the observed effects depend on acclimation time. For further details see *Supplementary Information*.

## Data analysis

The data were analysed using R (version 4.1.1) and RStudio using the packages lme4 (Bates et al. 2015), lmerTest (Kuznetsova et al. 2017), emmeans (Searle et al. 1980), car (Fox and Weisberg 2019) and plyr (Wickham 2011) for analyses and ggplot2, grid, cowplot, ggpubr and scales for plotting (Wickham 2009). Figures reporting mean values represent the least-squared means with 95% confidence level using a Tukey p-value adjustment for multiple comparisons unless stated otherwise.

### Metabolic density-dependence (Hypothesis 1)

For each species, we used the entire dataset to quantify the slope of the relationship between population metabolic rates (photosynthesis, post-illumination or dark respiration) and population biovolume (variables were log_10_ transformed). We performed this analysis separately for each metabolic rate and species using linear mixed effects models, including media treatment as a factor (with its interaction with population biovolume) and “flask ID” (parent population) as a random effect.

### Population biovolume growth

To describe the biovolume growth of each species over time and determine the growth phases (exponential phase and stationary phase), we fitted four types of growth models to the biovolume data of all replicates within a treatment. We chose this approach, rather than specifying a unique growth model, to allow for differences in growth patterns due to the media manipulations and species identity (Malerba et al. 2018). We selected the best fitting model based on the AIC, excluding the models that did not reach successful convergence. Model details are in the *Supplementary Information*.

### Metabolic responses to media treatments (Hypotheses 2 and 3)

Since all metabolic rates (except for *Dunaliella* post-illumination respiration) showed a different relationship with biovolume between exponential and stationary phase, we analysed these two phases separately to accurately determine the effects of media treatments. We considered biovolume 10 × 10^5^ µm^3^/µl as the cut-off based on the assessment of growth curves and the behaviour of the relationship between oxygen rates and biovolume (Figures S1-S2). Given the limited exposure time (3 hours), there was no effect of media treatments on the population biovolume of each species (Figures S1-S2; Tables S1-S4).

Each species and rate were analysed separately using linear mixed effects models that included population metabolic rate (photosynthesis, post illumination respiration, or dark respiration) as a function of population biovolume (covariate) and media treatment (predictor), and “flask_ID” (parent population) as a random effect. The initial models included an interaction between biovolume and media treatment. If this interaction was not significant, we compared models with and without interactions using the Akaike Information Criterion (AIC) and then fitted the data with the model that had the lowest AIC. Post-hoc comparisons were performed to test for differences in slopes (in case of a significant biovolume × media interaction) or intercepts between media treatments.

We excluded negative metabolic rates as they are biologically impossible, as well as day 0 because we did not measure on that day. Biovolume data were divided by 10^5^ before analyses to make the scales of the response and predictor variables more similar. Oxygen and biovolume data were log_10_-transformed in three cases to meet assumptions of normality (post-illumination respiration for *Dunaliella* and *Tetraselmis*; photosynthesis for *Tetraselmis*).

### Cell size plasticity

We tested for differences in cell size (μm^3^) in response to media manipulations using linear mixed effect models, with media treatment and experiment day as factorial predictors (since the relationship between cell volume and experiment day was non-linear). Flask_ID was again included as a random effect. As for metabolic rate analysis, day 0 was removed since we did not measure responses to media treatment on that day.

## Results

### H1: Metabolic density-dependence

We found density-dependent metabolic rates (slopes < 1) across most media conditions for all species (Figure 2), even when populations were resuspended in fresh media with plenty of nutrients. The only exception was for the dark respiration rates in artificial seawater (ASW), which showed isometric scaling for *Dunaliella* (at both incubation times) and hyper-allometric scaling for *Tetraselmis* (Table S5 for scaling exponents and confidence intervals). Below, we explore the effects of media treatments on the responses of each species since the relationship between metabolism and biovolume often varied between growth phases.

**Figure 2.**
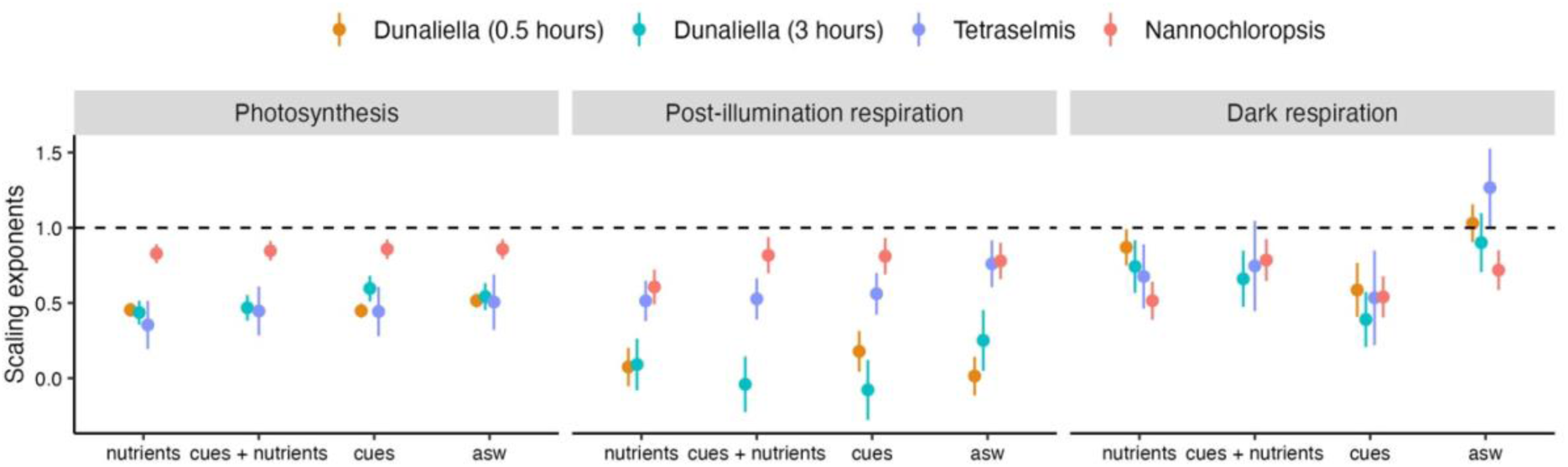
Scaling exponents (slopes of the log_10_-log_10_ data) of the relationship between population metabolic rates and population biovolume for each species and condition, throughout the experiment (not separated by growth phases, see H1). The dashed line indicates an exponent of 1 corresponding to a perfectly proportional (isometric) relationship between population metabolism and biovolume. Almost all species and conditions show values < 1, except dark respiration rates for *Dunaliella* and *Tetraselmis* in ASW.

### H2: Cues effects

The effects of media manipulations on metabolism were species-specific. Still, all three species showed some metabolic suppression in the presence of conspecific cues (Figure 3), for respiration rates more than photosynthesis although the strength of this reduction varied between species and growth phases (we delve on this below). Briefly, the addition of nutrients completely removed metabolic suppression in one species (*Tetraselmis*) but not in the other two (at least not for all metabolic rates; Figure 3).

**Figure 3.**
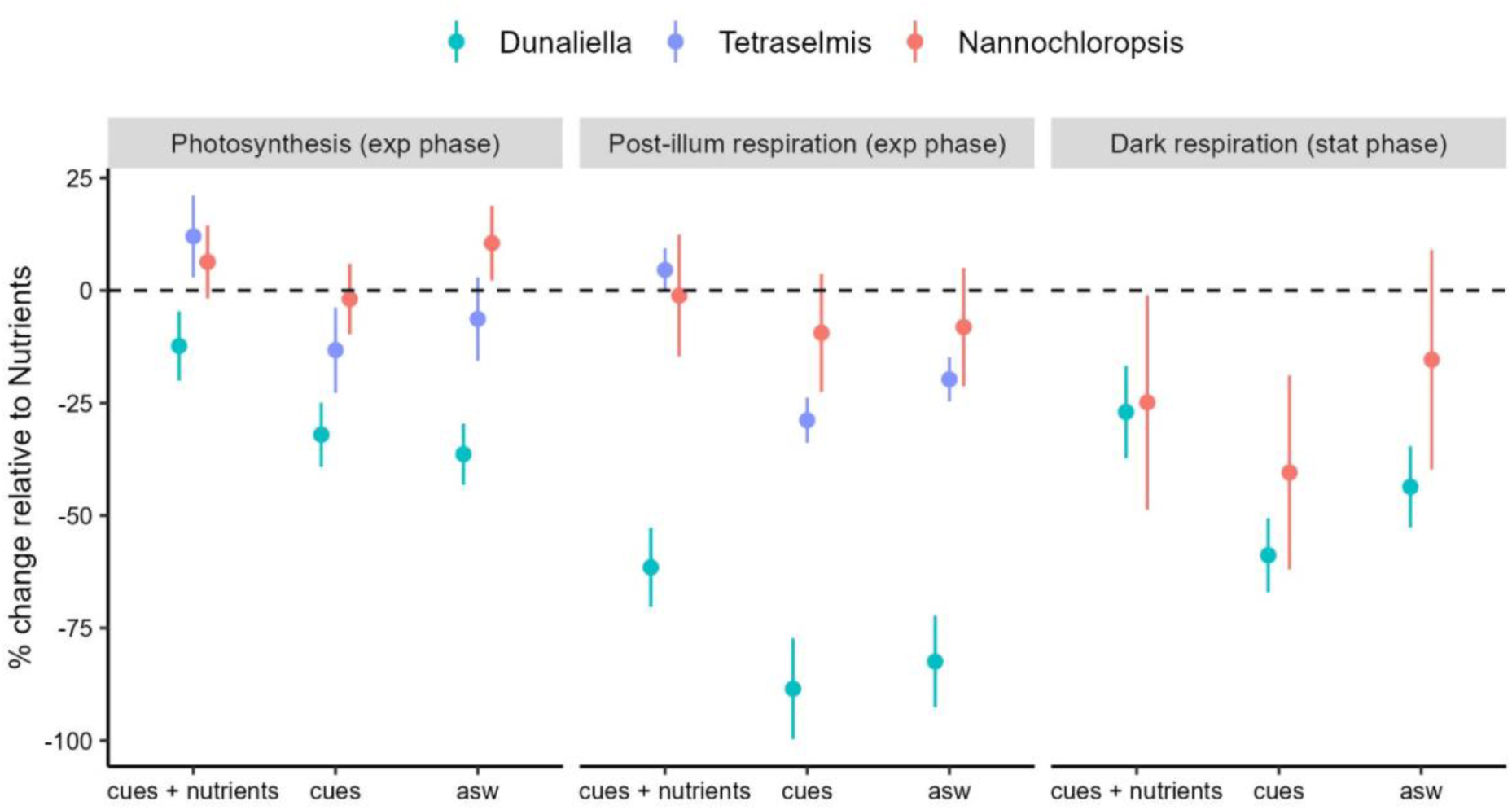
Percentage change of the intercept for each metabolic rate in each media treatment (“cues + nutrients”, “cues”, or “ASW”) relative to the “nutrients” treatment used as a reference (dashed line). Intercepts are the estimated marginal means at fixed biovolume calculated from post-hoc tests on the linear mixed models for each metabolic rate. A negative value means that metabolic rates were lower in that specific media treatment in comparison to those measured in the “nutrients” treatment.

*Dunaliella* showed consistent responses to both nutrients and cues for almost all metabolic rates and growth phases (Figure 4) in the direction we anticipated (Figure 1). Increasing exposure time from 0.5 to 3 hours strengthened responses (particularly to nutrients, Figure 4, Tables S6-S7), suggesting some delay in how cells adjusted their metabolic fluxes. In exponential phase, photosynthesis rates increased faster with biovolume in the treatments with nutrients (“nutrients” and “cues + nutrients”) than in artificial seawater (Figure 4a; biovolume × media: F_3, 142_ = 6.68, p = 0.0003; Table S7). But, across all metabolic rates, media manipulations had stronger effects on the intercepts (metabolic level; Figure 4) than on the slopes: populations had the highest photosynthesis and respiration rates when exposed to nutrients alone. The presence of cues reduced metabolic rates (sometimes more than artificial seawater); this reduction was weakened, but not eliminated, by the addition of nutrients (“cues + nutrients”). This trend was significant for photosynthesis during the exponential phase (Figure 4a), post-illumination respiration during the whole experiment (Figure 4c), and dark respiration in the stationary phase (Figure 4d, Tables S6-7). In stationary phase, photosynthesis showed the same pattern, but rates were significantly higher in the presence of nutrients alone while all other treatments did not differ (Figure 4b). We had a poor model fit for respiration rates in exponential phase but, overall, there were no strong effects of media treatments (Figure S3, Tables S6-S7).

**Figure 4.**
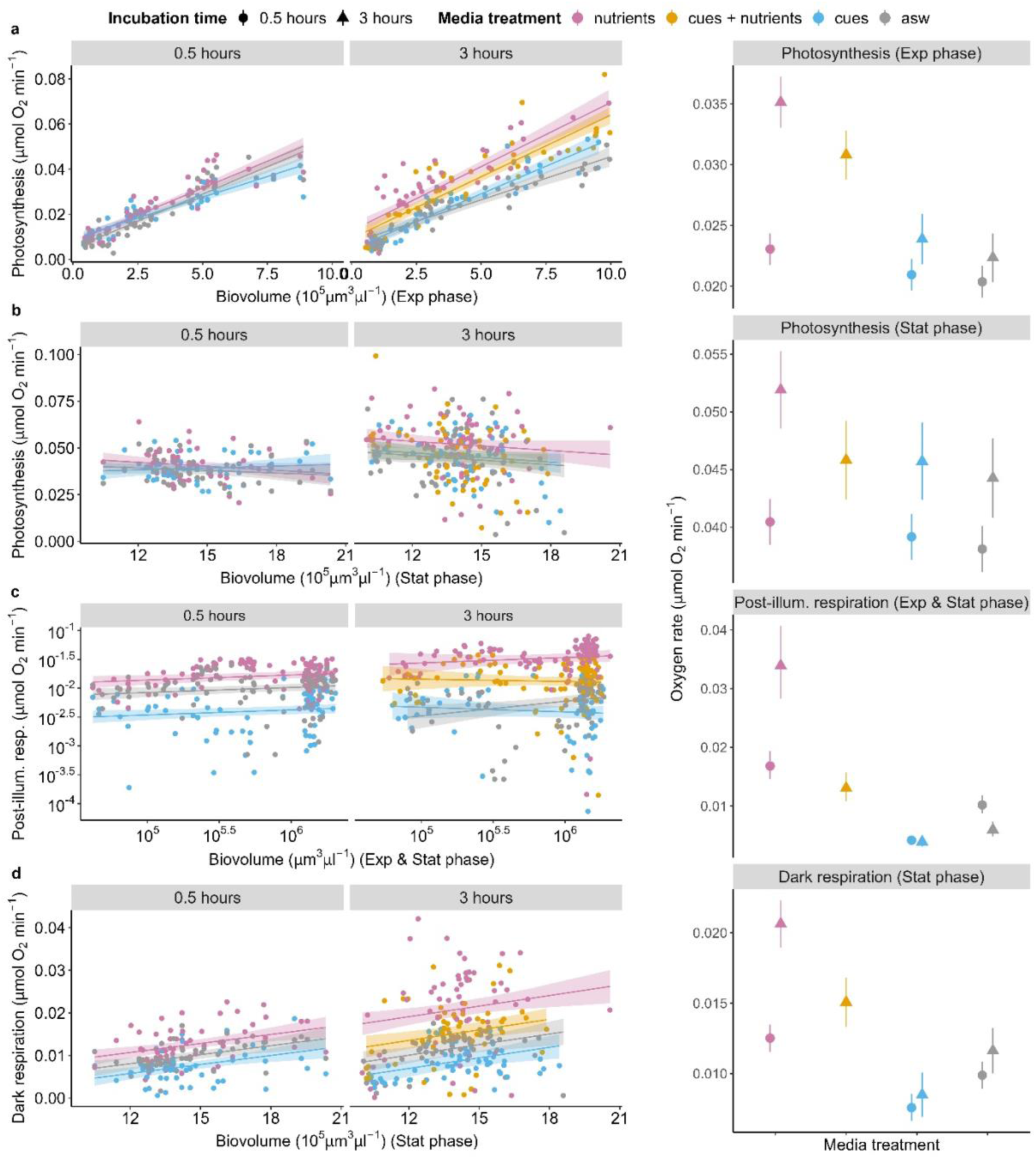
Changes in the metabolism of *Dunaliella tertiolecta* as a function of population biovolume for a) photosynthesis in exponential and b) stationary phase, c) post-illumination respiration across the entire biovolume range and d) dark respiration in stationary phase. The main effects of media manipulations on all these metabolic rates were on the intercepts (metabolic level, shown on the right column): rates were highest with nutrients alone and decreased with cues and artificial seawater; the addition of nutrients to cues weakened but did not eliminate metabolic suppression. In general, the strength of responses increased after 3 hours of exposure compared to 30 minutes.

The largest species, *Tetraselmis,* responded to nutrients but not to cues. Photosynthesis and post-illumination respiration showed similar patterns, but the effects were clearer in the latter: rates were higher when nutrients were added to the media, independently of cues (“nutrients” and “cues + nutrients” were identical), but differences between treatments weakened as biovolume increased due to a shallower slope in the treatments with nutrients (interaction media × biovolume; Figure 5a-b, Table S8). Dark respiration did not show a clear response to media manipulations except that “cues” had the shallowest slope (but they were significantly different to ASW only) (Figure 5c; Table S8).

**Figure 5.**
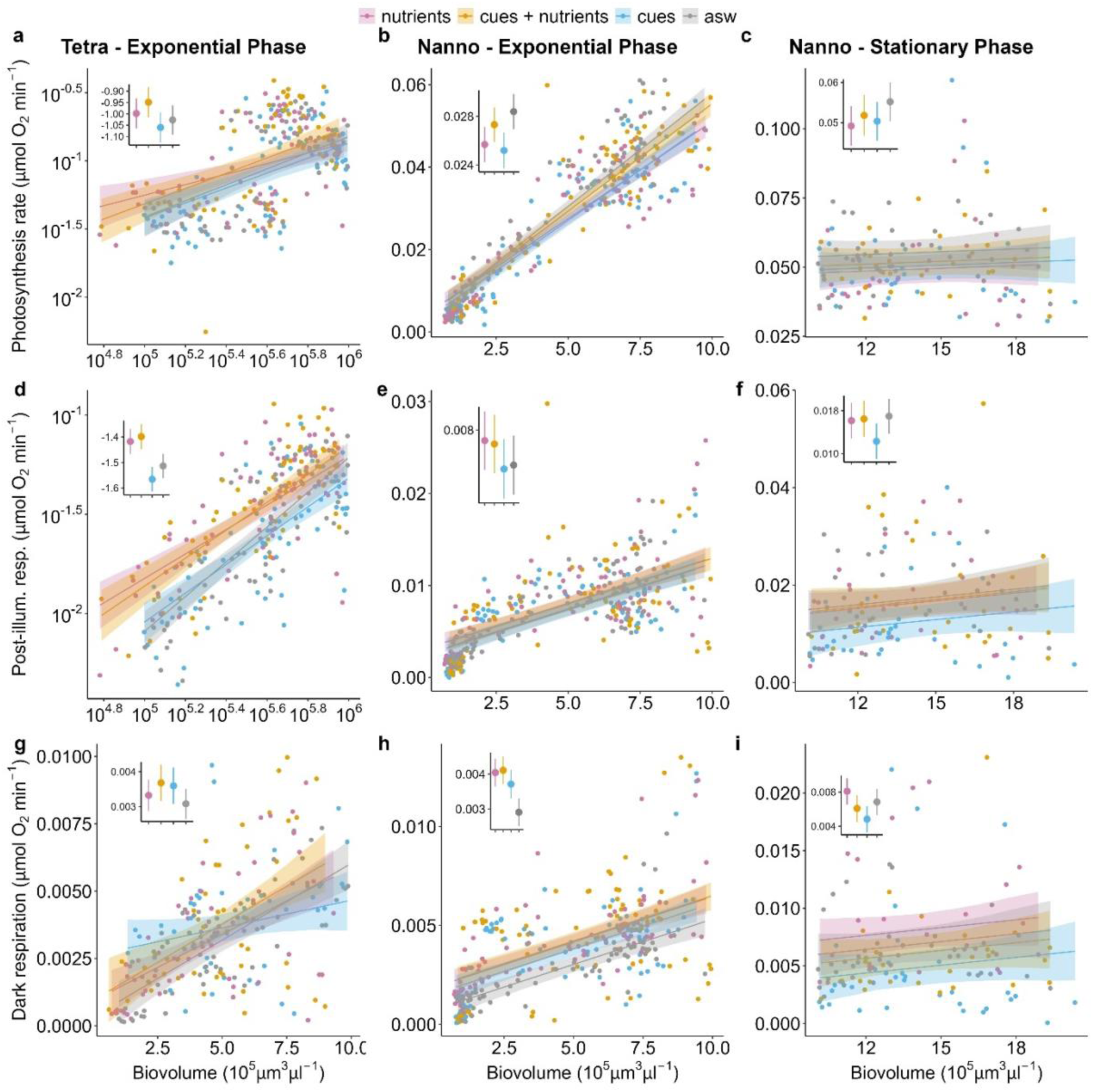
*Tetraselmis sp.* (a-c) and *Nannochloropsis granulata* (d-i) metabolic rates as a function of population biovolume during exponential phase (both species) and stationary phase (only for *Nannochloropsis*). The insert in each plot shows the intercepts for each media treatment (at fixed biovolume where there was an interaction). *Tetraselmis* photosynthesis and post-illumination respiration increased in response to nutrients, which completely removed the effect of cues (a-b). *Nannochloropsis* showed no or subtler responses to media manipulations for most rates and growth phases. The most notable effect was on dark respiration rates in stationary phase (i), where cues suppressed metabolism and the addition of nutrients decreased but did not remove this effect, as observed for *Dunaliella*. See Tables S8-S9 for analyses.

The smallest species, *Nannochloropsis,* had the weakest overall response to media manipulations, but cues still affected its respiration rates. This effect was particularly visible on dark respiration in stationary phase: respiration rates were highest with nutrients alone and lowest with cues (Figure 5i, Table S9). As for *Dunaliella*, the addition of nutrients to the cues reduced but did not negate metabolic suppression. Post-illumination respiration showed a similar but weaker pattern in stationary phase (Figure 5h). All other rates/growth phases showed negligible or non-significant effects (Table S9).

### H3: Cell size plasticity

While species size did not mediate metabolic responses in the way we anticipated (no support for H3), the cell size of each species responded to media manipulations within the 3-hour exposure, particularly in exponential phase. Within the first few days of exponential growth the three species showed a similar trend: cells were typically smaller when exposed to nutrients alone and larger when exposed to cues or ASW; the treatment “cues + nutrients” was often intermediate (Figure S4). Size differences between treatments were transient and often disappeared over the course of the experiment (Tables S10-S12), with only *Tetraselmis* showing consistent effects of media manipulations across days. Overall, all species showed remarkable cell size plasticity across growth phases and media treatments. The max. change in cell size was 71.4% for *Dunaliella*; 89.9% for *Tetraselmis*; 106.6% for *Nannochloropsis*. Changes in cell size were greater over time (i.e. between exponential and stationary phase) than in response to media manipulations within each day (Figure S4).

## Discussion

The fact that *per capita* metabolism declines in denser populations for many species is of biological significance because metabolism is linked to growth (DeLong et al. 2014). Reductions in resource availability are an obvious driver of metabolic suppression but not the only one (DeLong et al. 2014; Lovass et al. 2020). Metabolic suppression might also stem from active adjustments to the presence of conspecifics. Understanding the contribution of these two processes (resource limitation vs active adjustments) is important to forecast and model production. We explored the relative importance of nutrients and conspecific cues on three aspects of metabolism (photosynthesis, post-illumination, dark respiration) of three phytoplankton species. Overall we found that all metabolic rates are density-dependent even when populations are provided with plenty of fresh nutrients (H1). This density-dependence can in part be driven by light limitation as population biomass increases. However, light competition cannot explain the metabolic suppression we observed in response to cues when comparing populations at the same biomass density.

We hypothesised that organisms would reduce their metabolism in response to cues if these indicate crowding, and that metabolic suppression would be relieved by nutrient availability (H2). Our results were consistent with this hypothesis only for one species (*Dunaliella*). The smallest species (*Nannochloropsis*) showed a similar pattern but only for dark respiration and only in stationary phase. The largest species (*Tetraselmis*), instead, did not respond to cues but rather to nutrients (albeit we could not test responses in stationary phase). Therefore, differences in metabolic responses between species were not mediated by their cell size – at least not in the direction we anticipated (H3). In fact, the smallest species was the least sensitive to nutrients (*Nannochloropsis*) and the largest the most responsive (*Tetraselmis*).

This was contrary to our hypothesis because small cells have a higher surface-area to volume ratio and might be able to acquire nutrients more rapidly (Yoshiyama and A. Klausmeier 2008); in addition they have less capacity for storage (Malerba et al. 2018). Thus, we anticipated that the small *Nannochloropsis* might be more responsive to nutrients and cues if these signal crowding and a potential reduction in resource availability. A consideration is that we standardised populations by biovolume because total biovolume is the main driver of metabolism and growth in phytoplankton (Malerba et al. 2017; Fant and Ghedini 2024). This approach however meant that the small species was at high cell densities since the beginning of the experiment, which might have constrained its responses. Still, we think that this explanation is unlikely because *Nannochloropsis* showed metabolic suppression to cues in stationary phase, when cell density was highest. Sensitivity to nutrients and cues might thus be mediated by traits other than size but what these traits are is unclear.

Despite variation between species, metabolic rates were often reduced in the presence of cues, even during exponential phase when nutrients were not limiting in the spent media (since populations were still growing). This pattern (albeit of varying strength) suggests that organisms use conspecific cues to regulate their metabolic activity. Lowering metabolism might provide a fitness advantage by reducing resource requirements (Briddon et al. 2025; Auer, Salin, Rudolf, et al. 2015). The addition of nutrients led to a partial or complete recovery of metabolic rates, showing that organisms use cues in context: a surplus of nutrients can remove the metabolic suppression triggered by cues, since lowering metabolism has probably no fitness benefit in this context (Auer et al. 2020). Interestingly, we never observed an increase in metabolic rates in response to cues but only a decrease or no change.

Generally, photosynthesis was more affected by nutrients and less affected by cues than respiration. In particular, post-illumination respiration often tracked photosynthesis (as it reflects the metabolic costs during the light phase) (Beardall et al. 1994), but dark respiration and photosynthesis rates seemed to be differentially regulated. The downregulation of metabolism in response to cues (spent media) was stronger for dark respiration in stationary phase, possibly because nutrients were lowest and cell concentration highest. The addition of nutrients in this case did not recover “normal” respiration rates while it did for photosynthesis. Interestingly, respiration rates were often lower in spent media (which had both conspecific cues and the unused nutrients) than in artificial seawater (which had no nutrients at all). This result adds to the growing evidence that cells can independently adjust processes of energy production and metabolic expenditure, in response to changes in environmental conditions such as warming (Schaum et al. 2017; Padfield et al. 2016) and species interactions (Briddon et al. 2025). Maintaining energy production through photosynthesis (or feeding) while downregulating maintenance metabolism can be beneficial in crowded conditions, even if a lower respiration rate might over time reduce growth and reproduction (Burton et al. 2011; Ghedini et al. 2017).

We cannot exclude that metabolic suppression was caused by the properties of spent media (e.g. pH or waste products) (Narla et al. 2025). However photosynthesis and respiration rates were also downregulated in the first days of growth when waste products are low. More generally, we do not see strong interactions between media manipulations and biovolume, indicating that metabolic responses to nutrients and cues are consistent across biovolume densities and growth days.

The presence of nutrients and cues not only altered metabolism but also cell size. Size responses were somewhat disconnected from metabolic responses within species – but were quite consistent between species: cells rapidly (within 3 hours) decreased in volume when nutrients were added, and this size reduction was often limited by cues. This result suggests that nutrient limitation increases cell volume in agreement with some previous studies (Peter and Sommer 2013; Marañón 2015), even though others found an increase (Yan et al. 2021). Smaller cells have lower storage capacity than larger cells (Verdy et al. 2009) but have higher metabolic rates (Marañón 2015), light absorption (less package effect than in larger cells) and nutrient uptake ability per unit volume (Litchman et al. 2007; Edwards et al. 2011). Hence plastic reductions in cell size might be optimal if cells are acquiring nutrients in preparation to cell division, taking advantage of more favourable environmental conditions. Indeed, the reduction in size was more pronounced during exponential phase at least for two of the species. Why cues would reduce this size response is unclear but again it suggests that conspecific cues can induce a regulatory response that is not always fully eliminated by nutrients, similarly to what observed for metabolism.

In summary, intraspecific signalling can regulate phytoplankton metabolic activity in combination with resource availability. While the strength of responses varied between species and growth phases, metabolic suppression to conspecific cues was eliminated by nutrients in some but not all cases. Therefore, active metabolic adjustments to cues can be a source of density-dependence, reducing production even when resources are not limiting. Quantifying the importance of metabolic “self-regulation” would improve our capacity to predict population growth (Flux 2001; Fahimipour et al. 2025) and might help explain the similarity of growth patterns across species and systems that differ in resource availability (Hatton et al. 2015; Fant and Ghedini 2024). Our study shows that the interplay of nutrients and cues in regulating metabolism is complex and species-specific. Yet, accounting only for resource availability might provide an incomplete description of how populations grow.

## Acknowledgements

We thank you Lorenzo Fant for insightful discussions on the design of this experiment. We also thank José Marques for improving the ImageJ script for image processing. This work was supported by an ERC Starting Grant by the European Union to G.G. (META_FUN, 101116029).

## Author contributions

RE and GG designed the experiment. RE performed the experiments and analysed the data with feedback from GG and ALH. RE drafted the manuscript and all authors contributed to the final version.

