## Supplementary Information for "Drivers of metabolic density-dependence: how resource availability and conspecific cues affect phytoplankton metabolism"

### Supplementary Materials and Methods

*Additional experiment on Dunaliella tertiolecta*

For this experiment, we used a similar protocol to the one described in the main text, starting with 104 parent populations of *Dunaliella* grown in T25 cell culture flasks (Sarstedt, Germany), filled up to a final volume of 35 mL with ASW F medium. The initial biovolume of *Dunaliella* was the same as in the main experiments in each flask ($5 \times 10^4$ $\mu m^3/\mu l$).

Each sampling day, we used eight parent populations to generate, from each of them, three new populations in 10 mL falcon tubes resulting in 24 sample units per day. In addition, we sampled 1mL from each parent population flask which we stained with 1% Lugol's iodine for biovolume determination. The twenty-four 10 mL samples were centrifuged at 3500 rpm for 10 minutes and the algal pellets were resuspended in three different types of media: 1) "nutrients", 3) "cues", 4) "ASW" which are the same described above (Box 1), except for the treatment #2 ("cues + nutrients") which we did not have here.

After resuspension, the cultures were incubated for 30 minutes before measuring metabolic rates as described above. Metabolic rates were adjusted for the metabolism measured in blanks. We prepared four "control blanks" filled with spent media from the "cues" treatment and four "fresh media blanks" from the samples that were resuspended in fresh media ("nutrients" and "ASW") which had few to no bacteria. This process was repeated on thirteen occasions to cover the different growth phases.

*Population biovolume growth*

The models tested were 1) a logistic-type sinusoidal growth model with lower asymptote forced to 0 (three-parameter logistic curve); 2) a logistic-type sinusoidal growth model with non-zero lower asymptote (four-parameter logistic curve); 3) a Gompertz-type sinusoidal growth model (three-parameter Gompertz curve); 4) a modified Gompertz-type sinusoidal growth model including population decline after reaching a maximum (four-parameter Gompertz-like curve including mortality). We selected the best fitting model based on the AIC, excluding the models that did not reach successful convergence. Note that, for the 30 min incubation experiment of *Dunaliella*, we fitted growth models to the data collected from each parent flask (not from each media treatment). Due to the short exposure period, we assumed that morphological traits and growth patterns would be similar across treatments.

<u>Growth models abbreviations used in Tables S1-S4</u>

*Logistic type sinusoidal model parameters (logis):* Asym – asymptote; Xmid – inflection point; scal – scaling factor.

*Logistic type sinusoidal model with four parameters (fpl):* A – maximum asymptote; B – steepness of the curve; Xmid – inflection point; scal – scaling factor.

*Gompertz-type sinusoidal model parameters (Gomp):* Asym – asymptote; B2 – initial growth rate; B3 – rate of decay of growth.

*Modified Gompertz-type sinusoidal model parameters (Modl)*: ymax – maximum growth value asymptotically; umin – growth rate; tmax – time reference; k – rate of deceleration.

### Supplementary Figures and Tables

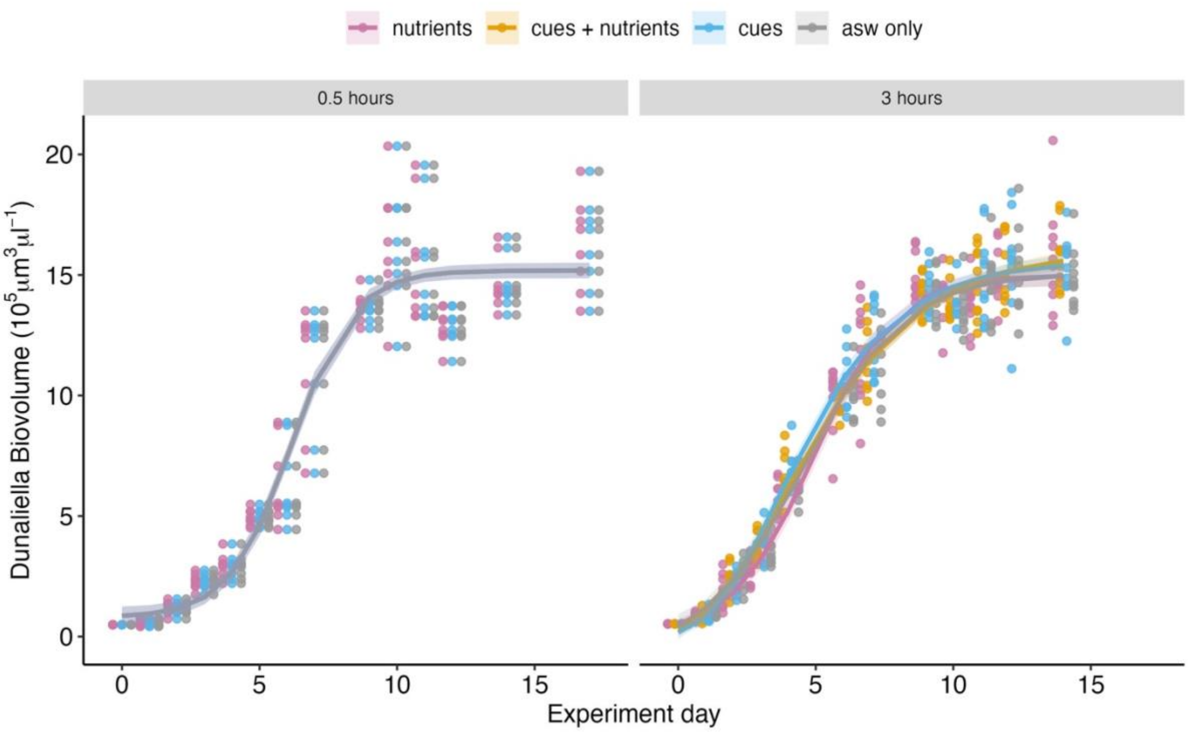


**Figure S1**. Increases in *Dunaliella* biovolume over time (experiment day). Growth patterns are similar between media treatments (colours) and exposure times (panel grids). Each experiment day represents a new parent population. Solid lines present fitted growth models (see Table S1 and S2 for growth model selection). Note that models overlap in left panel because samples were collected on the parent population and not on individual samples, given the short exposure time (0.5 hours).

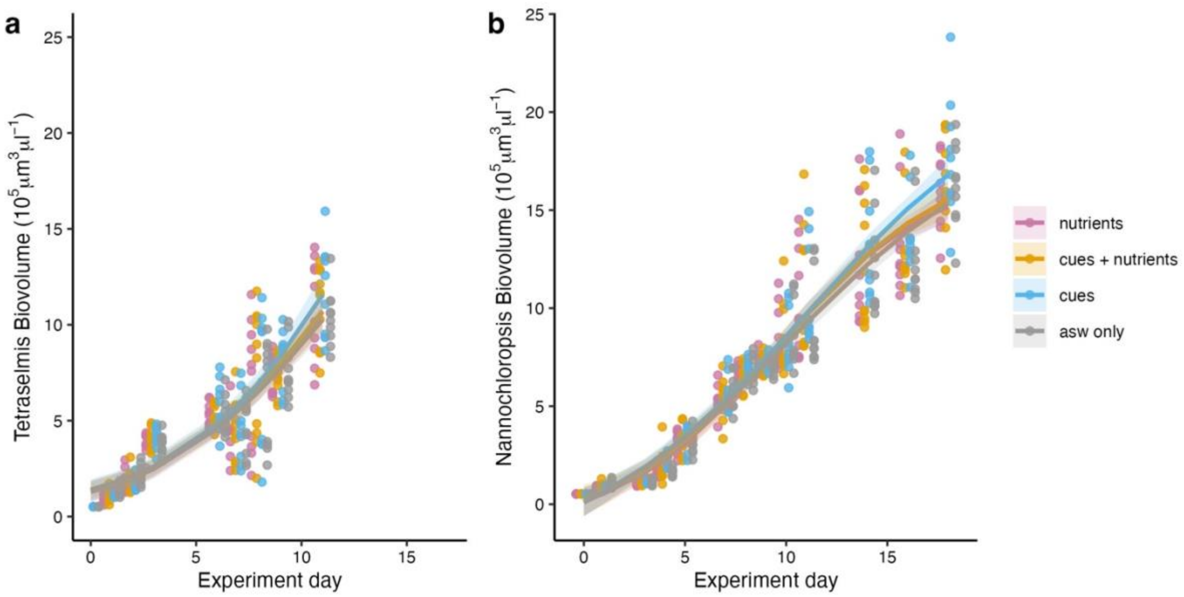


**Figure S2**. Increases in *Tetraselmis* (a) and *Nannochloropsis* (b) biovolume over time during the experiment. For *Tetraselmis* we collected data only in the exponential phase because culture became contaminated. Growth patterns are similar between media treatments, indicating no population growth over the 3 hour exposure. Solid lines present fitted growth models (see Table S3 and S4 for growth model selection).

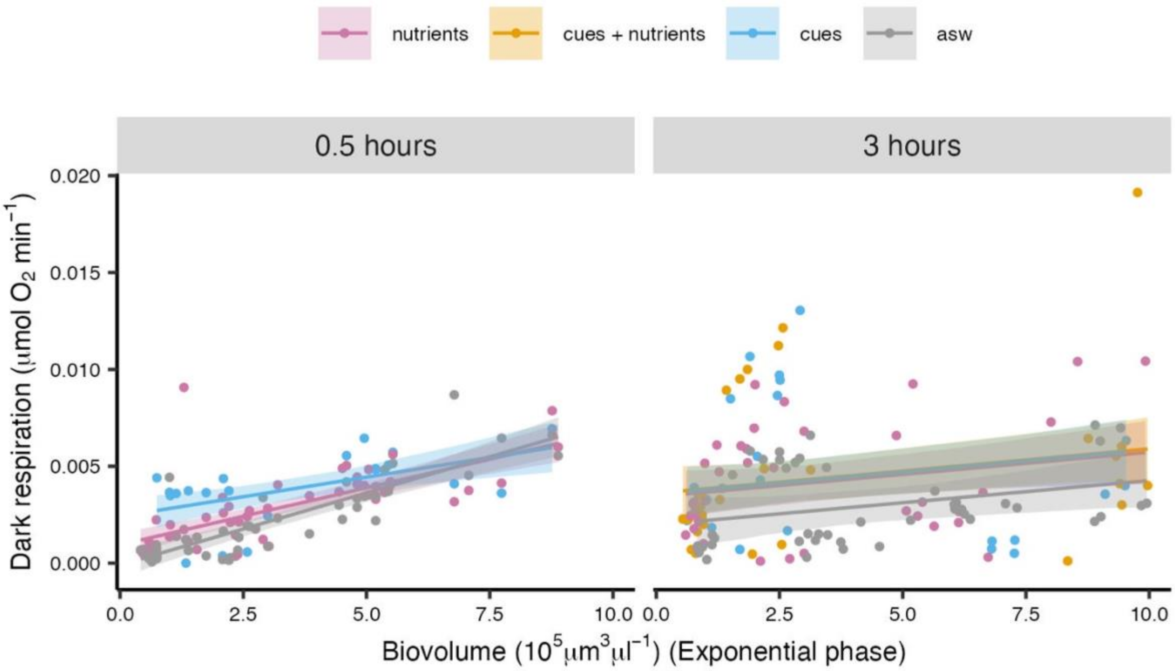


**Figure S3**. *Dunaliella* dark respiration rate during exponential phase at the different exposure times (panel grids) shows no clear effects of media treatments.

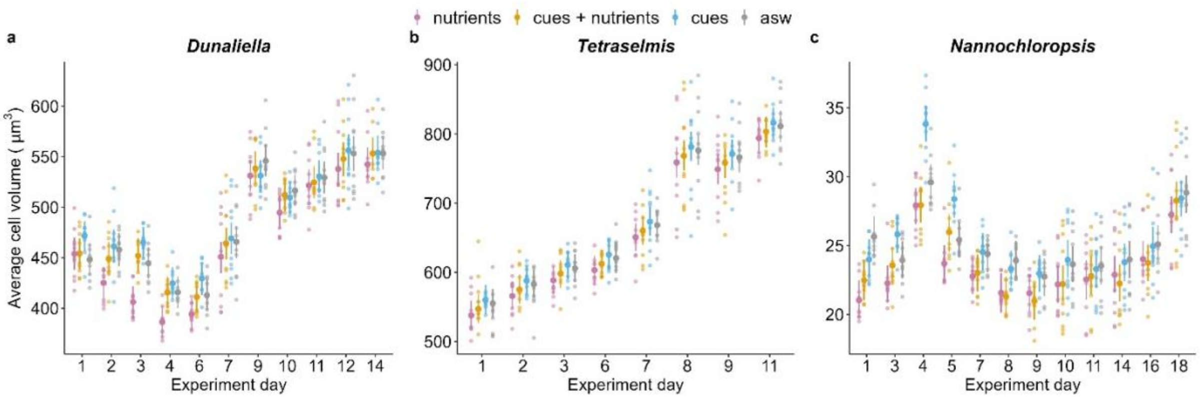


**Figure S4.** The average cell volume of each species changed over time (cells tended to be larger in stationary phase) but, within each day, cell size also responded to media manipulations. The effect of media treatments was variable between species and days (see Tables S10-S12) but, in exponential phase, cells were generally smaller when exposed to nutrients alone; cues or ASW tended to reduce this size response.

**Table S1.** Growth models fitted to *Dunaliella tertiolecta* biovolume (µm$^3$/µl) (0.5 hours exposure). The best fitting model (Fpl) was selected based on the Akaike information criterion (AIC), excluding the models that did not reach successful convergence.

| Biovolume growth models | | | | |
|---|---|---|---|---|
| Model | | df | AIC | |
| Logis | | 4 | 9032.293 | |
| Fpl | | 5 | 9023.892 | |
| Gomp | | 4 | 9073.697 | |
| Modl | | 5 | 9072.646 | |
| Best fitting model - fpl | | | | |
| Parameters | Estimate | Standard Error | T value | Pr (> \|t\|) |
| A | $7.99\times10^{4}$ | $2.14\times10^{4}$ | 3.739 | 0.0002 (***) |
| B | $1.52\times10^{6}$ | $1.65\times10^{4}$ | 92.256 | $<2\times10^{-16}$ (***) |
| Xmid | 6.17 | $8.42\times10^{-2}$ | 73.309 | $<2\times10^{-16}$ (***) |
| scal | 1.14 | $8.40\times10^{-2}$ | 13.522 | $<2\times10^{-16}$ (***) |

**Table S2.** Growth models fitted to *Dunaliella tertiolecta* biovolume data ($\mu m^3/\mu l$) collected for each treatment (3 hours exposure; 4 treatments). The best fitting model was selected based on the Akaike information criterion (AIC), excluding the models that did not reach successful convergence.

| **Biovolume growth models - Nutrients** | | | | |
|---|---|---|---|---|
| Model | | df | AIC | |
| Logis | | 4 | 3159.418 | |
| Fpl | | 5 | 3161.197 | |
| Gomp | | 4 | 3162.674 | |
| Modl | | 5 | 3164.618 | |
| Best fitting model - Logis | | | | |
| Parameters | Estimate | Standard Error | T value | Pr (> \|t\|) |
| Asym | $1.50\times10^{6}$ | $2.14\times10^{4}$ | 69.15 | $<2\times10^{-16}$ (***) |
| Xmid | 4.93 | 0.116 | 42.43 | $<2\times10^{-16}$ (***) |
| scal | 1.48 | $8.84\times10^{-2}$ | 16.75 | $<2\times10^{-16}$ (***) |
| **Biovolume growth models – Nutrients + Cues** | | | | |
| Model | | df | AIC | |
| Logis | | 4 | 3107.494 | |
| Fpl | | 5 | 3099.138 | |
| Gomp | | 4 | 3093.173 | |
| Best fitting model - Gomp | | | | |
| Parameters | Estimate | Standard Error | T value | Pr (> \|t\|) |
| Asym | $1.60\times10^{6}$ | $2.62\times10^{4}$ | 61.08 | $<2\times10^{-16}$ (***) |
| B2 | 3.97 | 0.25 | 15.88 | $<2\times10^{-16}$ (***) |
| B3 | 0.7 | $1.28\times10^{-2}$ | 54.68 | $<2\times10^{-16}$ (***) |
| **Biovolume growth models – Cues** | | | | |
| Model | | df | AIC | |
| Logis | | 4 | 3147.821 | |
| Fpl | | 5 | 3145.58 | |
| Gomp | | 4 | 3141.122 | |
| Modl | | 5 | 3143.121 | |
| Best fitting model - Gomp | | | | |
| Parameters | Estimate | Standard Error | T value | Pr (> \|t\|) |
| Asym | $1.56\times10^{6}$ | $2.64\times10^{4}$ | 59.12 | $<2\times10^{-16}$ (***) |
| B2 | 4.46 | 0.392 | 11.39 | $<2\times10^{-16}$ (***) |
| B3 | 0.663 | $1.69\times10^{-2}$ | 39.21 | $<2\times10^{-16}$ (***) |
| **Biovolume growth models – ASW** | | | | |
| Model | | df | AIC | |
| Logis | | 4 | 3121.681 | |
| Fpl | | 5 | 3120.887 | |
| Gomp | | 4 | 3120.967 | |
| Modl | | 5 | 3121.076 | |
| Best fitting model - Fpl | | | | |
| Parameters | Estimate | Standard Error | T value | Pr (> \|t\|) |
| A | $-8.66\times10^{4}$ | $6.01\times10^{4}$ | -1.44 | 0.153 |
| B | $1.51\times10^{6}$ | $2.37\times10^{4}$ | 63.58 | $<2\times10^{-16}$ (***) |
| Xmid | 4.56 | 0.192 | 23.84 | $<2\times10^{-16}$ (***) |
| scal | 1.86 | 0.166 | 11.22 | $<2\times10^{-16}$ (***) |

**Table S3.** Growth models fitted to *Tetraselmis sp.* biovolume data (μm$^3$/μl) collected for each treatment (3 hours exposure; 4 treatments; data until day 11). The only model that consistently fitted the data was Logis – logistic type sinusoidal growth model with lower asymptote forced to zero.

| **Biovolume growth models - Nutrients** | | | | |
|---|---|---|---|---|
| Best fitting model – Logis | | | | |
| Parameters | Estimate | Standard Error | T value | Pr (> \|t\|) |
| Asym | $2.58 \times 10^{6}$ | $1.63 \times 10^{6}$ | 1.587 | 0.1161 |
| Xmid | 12.5 | 4.66 | 2.682 | 0.0088 (**) |
| scal | 4.24 | 0.92 | 4,749 | $7.99 \times 10^{-6}$ (***) |
| **Biovolume growth models – Nutrients + Cues** | | | | |
| Best fitting model – Logis | | | | |
| Parameters | Estimate | Standard Error | T value | Pr (> \|t\|) |
| Asym | $2.59 \times 10^{6}$ | $1.64 \times 10^{6}$ | 1.577 | 0.1184 |
| Xmid | 12.5 | 4.76 | 2.621 | 0.0103 (*) |
| Scal | 4.3 | 0.917 | 4.69 | $1.01 \times 10^{-5}$ (***) |
| **Biovolume growth models – Cues** | | | | |
| Best fitting model – Logis | | | | |
| Parameters | Estimate | Standard Error | T value | Pr (> \|t\|) |
| Asym | $3.54 \times 10^{6}$ | $3.27 \times 10^{6}$ | 1.082 | 0.2824 |
| Xmid | 14.2 | 6.61 | 2.149 | 0.0344 (*) |
| Scal | 4.43 | 0.998 | 4.445 | $2.58 \times 10^{-5}$ (***) |
| **Biovolume growth models – ASW** | | | | |
| Best fitting model – Logis | | | | |
| Parameters | Estimate | Standard Error | T value | Pr (> \|t\|) |
| Asym | $2.49 \times 10^{6}$ | $1.44 \times 10^{6}$ | 1.737 | 0.086 (.) |
| Xmid | 12.6 | 4.48 | 2.811 | 0.0061 (**) |
| scal | 4.47 | 0.858 | 5.209 | $1.26 \times 10^{-6}$ (***) |

**Table S4.** Growth models fitted to *Nannochloropsis granulata* biovolume data ($\mu m^3/\mu l$) collected for each treatment (3 hours exposure; 4 treatments). Only two growth models fitted the growth data (logs, fpl). The best fitting model was selected based on the Akaike information criterion (AIC), excluding the models that did not reach successful convergence.

| | | | | |
|---|---|---|---|---|
| **Biovolume growth models - Nutrients** | | | | |
| Model | | df | AIC | |
| Logis | | 4 | 3489.283 | |
| Fpl | | 5 | 3488.499 | |
| Best fitting model – Fpl | | | | |
| Parameters | Estimate | Standard Error | T value | Pr (> \|t\|) |
| A | $-1.34\times10^{5}$ | $1.08\times10^{5}$ | -1.248 | 0.214 |
| B | $1.71\times10^{6}$ | $1.31\times10^{5}$ | 13.011 | $<2\times10^{-16}$ (***) |
| Xmid | 9.45 | 0.501 | 18.859 | $<2\times10^{-16}$ (***) |
| scal | 3.90 | 0.751 | 5.197 | $7.94\times10^{-7}$ (***) |
| **Biovolume growth models – Nutrients + Cues** | | | | |
| Model | | df | AIC | |
| Logis | | 4 | 3516.165 | |
| Fpl | | 5 | 3515.632 | |
| Best fitting model – Fpl | | | | |
| Parameters | Estimate | Standard Error | T value | Pr (> \|t\|) |
| A | $-1.62\times10^{5}$ | $1.40\times10^{5}$ | -1.156 | 0.25 |
| B | $1.80\times10^{6}$ | $1.86\times10^{5}$ | 9.685 | $<2\times10^{-16}$ (***) |
| Xmid | 9.72 | 0.654 | 14.869 | $<2\times10^{-16}$ (***) |
| scal | 4.26 | 0.997 | 4.274 | $3.75\times10^{-5}$ (***) |
| **Biovolume growth models - Cues** | | | | |
| Model | | df | AIC | |
| Logis | | 4 | 3496.853 | |
| Fpl | | 5 | 3492.978 | |
| Best fitting model – Fpl | | | | |
| Parameters | Estimate | Standard Error | T value | Pr (> \|t\|) |
| A | $-2.85\times10^{5}$ | $2.04\times10^{5}$ | -1.401 | 0.1636 |
| B | $2.22\times10^{6}$ | $3.78\times10^{5}$ | 5.874 | $3.55\times10^{-8}$ (***) |
| Xmid | 11 | 1.16 | 9.448 | $2.42\times10^{-16}$ (***) |
| Scal | 5.47 | 1.54 | 3.558 | 0.0005 (***) |
| **Biovolume growth models – ASW** | | | | |
| Model | | df | AIC | |
| Logis | | 4 | 3481.365 | |
| Fpl | | 5 | 3476.678 | |
| Best fitting model - Fpl | | | | |
| Parameters | Estimate | Standard Error | T value | Pr (> \|t\|) |
| A | $-2.97\times10^{5}$ | $2.11\times10^{5}$ | -1.406 | 0.1623 |
| B | $1.97\times10^{6}$ | $3.06\times10^{5}$ | 6.419 | $2.54\times10^{-9}$ (***) |
| Xmid | 10.1 | 0.943 | 10.732 | $<2\times10^{-16}$ (***) |
| scal | 5.45 | 1.56 | 3.5 | 0.0006 (***) |

**Table S5.** Linear mixed effects models quantifying the slope of the relationship between population metabolic rates (μmol $O_2$ $min^{-1}$) and population biovolume ($\mu m^3/\mu l$) in the different media treatments for all species (Resuspended media, 3 levels for 0.5 hours of exposure and 4 levels for 3 hours of exposure). Variables were $log_{10}$ transformed. "Flask_ID" identifying the parent population was included as a random effect in the model. CL = 95% confidence level.

| ***Dunaliella tertiolecta* Photosynthesis – 0.5 hours** | | | | | |
|---|---|---|---|---|---|
| Resuspended media | Slope | SE | df | Lower CL | Upper CL |
| Nutrients | 0.454 | 0.0235 | 204 | 0.408 | 0.5 |
| Cues | 0.448 | 0.0235 | 204 | 0.402 | 0.495 |
| ASW | 0.516 | 0.0235 | 204 | 0.47 | 0.562 |
| ***Dunaliella tertiolecta* Post-Illumination – 0.5 hours** | | | | | |
| Resuspended media | Slope | SE | df | Lower CL | Upper CL |
| Nutrients | 0.0745 | 0.065 | 272 | -0.0534 | 0.202 |
| Cues | 0.1781 | 0.0688 | 272 | 0.0427 | 0.314 |
| ASW | 0.0139 | 0.0652 | 272 | -0.1145 | 0.142 |
| ***Dunaliella tertiolecta* Dark Respiration – 0.5 hours** | | | | | |
| Resuspended media | Slope | SE | df | Lower CL | Upper CL |
| Nutrients | 0.87 | 0.0612 | 265 | 0.749 | 0.99 |
| Cues | 0.586 | 0.0912 | 275 | 0.407 | 0.766 |
| ASW | 1.029 | 0.0639 | 268 | 0.903 | 1.155 |
| ***Dunaliella tertiolecta* Photosynthesis – 3 hours** | | | | | |
| Resuspended media | Slope | SE | df | Lower CL | Upper CL |
| Nutrients | 0.436 | 0.0404 | 214 | 0.356 | 0.515 |
| Cues + Nutrients | 0.468 | 0.0431 | 214 | 0.382 | 0.553 |
| Cues | 0.596 | 0.0433 | 214 | 0.51 | 0.681 |
| ASW | 0.542 | 0.00455 | 214 | 0.453 | 0.632 |
| ***Dunaliella tertiolecta* Post Illumination – 3 hours** | | | | | |
| Resuspended media | Slope | SE | df | Lower CL | Upper CL |
| Nutrients | 0.0911 | 0.0874 | 347 | -0.0808 | 0.263 |
| Cues + Nutrients | -0.0406 | 0.094 | 347 | -0.2255 | 0.144 |
| Cues | -0.0777 | 0.101 | 347 | -0.2767 | 0.121 |
| ASW | 0.2509 | 0.103 | 347 | 0.049 | 0.453 |
| ***Dunaliella tertiolecta* Dark respiration – 3 hours** | | | | | |
| Resuspended media | Slope | SE | df | Lower CL | Upper CL |
| Nutrients | 0.742 | 0.0888 | 294 | 0.567 | 0.917 |
| Cues + Nutrients | 0.66 | 0.0948 | 296 | 0.474 | 0.847 |
| Cues | 0.39 | 0.0931 | 286 | 0.207 | 0.573 |
| ASW | 0.901 | 0.0995 | 291 | 0.705 | 1.097 |
| ***Tetraselmis sp.* Photosynthesis – 3 hours** | | | | | |
| Resuspended media | Slope | SE | df | Lower CL | Upper CL |
| Nutrients | 0.354 | 0.0811 | 208 | 0.194 | 0.514 |
| Cues + Nutrients | 0.446 | 0.0824 | 209 | 0.284 | 0.609 |
| Cues | 0.443 | 0.0832 | 208 | 0.279 | 0.607 |
| ASW | 0.506 | 0.0936 | 208 | 0.321 | 0.69 |
| ***Tetraselmis sp.* Post Illumination – 3 hours** | | | | | |
| Resuspended media | Slope | SE | df | Lower CL | Upper CL |
| Nutrients | 0.514 | 0.0683 | 208 | 0.379 | 0.648 |
| Cues + Nutrients | 0.527 | 0.0695 | 209 | 0.39 | 0.664 |
| Cues | 0.561 | 0.0702 | 209 | 0.422 | 0.699 |
| ASW | 0.76 | 0.0789 | 209 | 0.605 | 0.916 |
| ***Tetraselmis sp.* Dark respiration – 3 hours** | | | | | |
| Resuspended media | Slope | SE | df | Lower CL | Upper CL |
| Nutrients | 0.676 | 0.108 | 232 | 0.463 | 0.889 |
| Cues + Nutrients | 0.746 | 0.152 | 240 | 0.446 | 1.046 |
| Cues | 0.534 | 0.16 | 240 | 0.219 | 0.848 |
| ASW | 1.266 | 0.132 | 234 | 1.006 | 1.526 |

| *Nannochloropsis granulata* **Photosynthesis – 3 hours** | | | | | |
|---|---|---|---|---|---|
| Resuspended media | Slope | SE | df | Lower CL | Upper CL |
| Nutrients | 0.828 | 0.0322 | 245 | 0.764 | 0.891 |
| Cues + Nutrients | 0.846 | 0.0332 | 245 | 0.781 | 0.912 |
| Cues | 0.858 | 0.0332 | 245 | 0.792 | 0.923 |
| ASW | 0.857 | 0.0337 | 245 | 0.791 | 0.923 |
| *Nannochloropsis granulata* **Post Illumination – 3 hours** | | | | | |
| Resuspended media | Slope | SE | df | Lower CL | Upper CL |
| Nutrients | 0.606 | 0.0585 | 332 | 0.491 | 0.721 |
| Cues + Nutrients | 0.817 | 0.0611 | 339 | 0.697 | 0.937 |
| Cues | 0.81 | 0.0618 | 346 | 0.689 | 0.932 |
| ASW | 0.779 | 0.0614 | 333 | 0.658 | 0.9 |
| *Nannochloropsis granulata* **Dark respiration – 3 hours** | | | | | |
| Resuspended media | Slope | SE | df | Lower CL | Upper CL |
| Nutrients | 0.515 | 0.064 | 352 | 0.389 | 0.641 |
| Cues + Nutrients | 0.785 | 0.0706 | 389 | 0.646 | 0.924 |
| Cues | 0.54 | 0.0697 | 380 | 0.403 | 0.677 |
| ASW | 0.718 | 0.0672 | 353 | 0.586 | 0.85 |

**Table S6.** Linear mixed effect models and post-hoc comparisons testing the relationship between population metabolic rates (µmol $O_2$ $min^{-1}$) and population biovolume (µ$m^3$/µl) in the different media treatments (Resuspended media, 3 levels) for the algal species *Dunaliella tertiolecta* (0.5 hours exposure) in different growth phases. "Flask_ID" identifying the parent population was included as a random effect in the model. Interactions were removed based on AIC evaluation of the models. Each metabolic rate (photosynthesis, post-illumination and dark respirations) was analyzed independently. Biov = Biovolume. EMM = estimated marginal means. CL = 95% confidence level.

| **Photosynthesis Exponential phase** | | | | | | |
|---|---|---|---|---|---|---|
| | Sum Sq | Mean Sq | NumDF | DenDF | F value | Pr (>F) |
| Biov | 0.0038593 | 0.0038593 | 1 | 48 | 352.3475 | $< 2.20 \times 10^{-16}$ (***) |
| Resuspended media | 0.0001584 | 0.0000792 | 2 | 96 | 7.2304 | 0.0012 (**) |
| Biov. × Resuspended media | 0.002202 | 0.0001101 | 2 | 96 | 10.0533 | 0.0001 (***) |
| Post hoc test on biovolume × resuspended media | | | | | | |
| Resuspended media | Slope | SE | df | Lower CL | | Upper CL |
| Nutrients | 0.00479 | 0.000291 | 97.2 | 0.00421 | | 0.00537 |
| Cues | 0.00367 | 0.000291 | 97.2 | 0.0031 | | 0.00425 |
| ASW | 0.00483 | 0.000291 | 97.2 | 0.00426 | | 0.00541 |
| Contrasts | Estimate | SE | df | t.ratio | | p.value |
| Nutrients – Cues | $1.12 \times 10^{-3}$ | 0.000293 | 96 | 3.807 | | 0.0007 |
| Nutrients – ASW | $-4.34 \times 10^{-5}$ | 0.000293 | 96 | -0.148 | | 0.988 |
| Cues – ASW | $-1.16 \times 10^{-3}$ | 0.000293 | 96 | -3.955 | | 0.0004 |
| Post hoc test on resuspended media controlling for biovolume (= $3.2 \times 10^5$ µ$m^3$/µl) | | | | | | |
| Resuspended media | EMM | SE | df | Lower CL | | Upper CL |
| Nutrients | 0.0231 | 0.000656 | 97.2 | 0.0218 | | 0.0244 |
| Cues | 0.0209 | 0.000656 | 97.2 | 0.0196 | | 0.0223 |
| ASW | 0.0204 | 0.000656 | 97.2 | 0.0191 | | 0.0217 |
| Contrasts | Estimate | SE | df | t.ratio | | p.value |
| Nutrients – Cues | 0.00211 | 0.000662 | 96 | 3.188 | | 0.0054 |
| Nutrients – ASW | 0.002693 | 0.000662 | 96 | 4.068 | | 0.0003 |
| Cues – ASW | 0.000583 | 0.000662 | 96 | 0.88 | | 0.6541 |
| **Photosynthesis Stationary phase** | | | | | | |
| | Sum Sq | Mean Sq | NumDF | DenDF | F value | Pr (>F) |
| Biov | $2.64 \times 10^{-5}$ | $2.64 \times 10^{-5}$ | 1 | 52 | 0.6744 | 0.4153 |
| Resuspended media | $1.74 \times 10^{-4}$ | $8.71 \times 10^{-5}$ | 2 | 104 | 2.2283 | 0.1128 |
| Biov × Resuspended media | $1.62 \times 10^{-4}$ | $8.10 \times 10^{-5}$ | 2 | 104 | 2.0718 | 0.1311 |
| **Post-Illumination Exponential and Stationary phase** | | | | | | |
| | Sum Sq | Mean Sq | NumDF | DenDF | F value | Pr (>F) |
| $Log_{10}$(Biov) | 0.4431 | 0.4431 | 1 | 87.732 | 4.7078 | 0.0327 (*) |
| Resuspended media | 16.0132 | 8.0066 | 2 | 187.135 | 85.0656 | $< 2 \times 10^{-16}$ (***) |
| Post hoc test on resuspended media | | | | | | |
| Resuspended media | EMM | SE | df | Lower CL | | Upper CL |
| Nutrients | -1.77 | 0.0315 | 270 | -1.84 | | -1.71 |
| Cues | -2.38 | 0.0367 | 272 | -2.45 | | -2.31 |
| ASW | -1.99 | 0.0322 | 271 | -2.06 | | -1.93 |
| Contrasts | Estimate | SE | df | t.ratio | | p.value |
| Nutrients – Cues | 0.605 | 0.0466 | 196 | 12.989 | | <.0001 |
| Nutrients – ASW | 0.217 | 0.0432 | 180 | 5.034 | | <.0001 |
| Cues – ASW | -0.387 | 0.0469 | 194 | -8.252 | | <.0001 |

| Dark Respiration Exponential phase | | | | | | |
|---|---|---|---|---|---|---|
| | Sum Sq | Mean Sq | NumDF | DenDF | F value | Pr (>F) |
| Biov | $1.65 \times 10^{-4}$ | $1.65 \times 10^{-4}$ | 1 | 47.5 | 101.1287 | $2.37 \times 10^{-13}$ (***) |
| Resuspended media | $2.85 \times 10^{-5}$ | $1.43 \times 10^{-5}$ | 2 | 81.391 | 8.7703 | 0.0003 (***) |
| Biov × Resuspended media | $9.25 \times 10^{-6}$ | $4.62 \times 10^{-6}$ | 2 | 78.415 | 2.8407 | 0.0644 (.) |
| Post hoc test on biovolume × resuspended media (p = 0.06) | | | | | | |
| Resuspended media | Slope | SE | df | Lower CL | Upper CL | |
| Nutrients | 0.000588 | $8.19 \times 10^{-5}$ | 113 | 0.000425 | 0.00075 | |
| Cues | 0.000408 | $1.21 \times 10^{-4}$ | 113 | 0.000168 | 0.000647 | |
| ASW | 0.000744 | $8.41 \times 10^{-5}$ | 113 | 0.000578 | 0.000911 | |
| Contrasts | Estimate | SE | df | t.ratio | p.value | |
| Nutrients – Cues | 0.00018 | 0.000144 | 86.7 | 1.252 | 0.4264 | |
| Nutrients – ASW | -0.000157 | 0.000115 | 68.5 | -1.368 | 0.3634 | |
| Cues – ASW | -0.000337 | 0.000145 | 87.3 | -2.322 | 0.058 | |
| Post hoc test on resuspended media controlling for biovolume (= $3.32 \times 10^5$ µm³/µl) | | | | | | |
| Resuspended media | EMM | SE | df | Lower CL | Upper CL | |
| Nutrients | 0.00291 | 0.000185 | 113 | 0.00254 | 0.00327 | |
| Cues | 0.00376 | 0.000284 | 113 | 0.00319 | 0.00432 | |
| ASW | 0.00238 | 0.000191 | 113 | 0.002 | 0.00275 | |
| Contrasts | Estimate | SE | df | t.ratio | p.value | |
| Nutrients – Cues | -0.000851 | 0.000334 | 88.5 | -2.549 | 0.0332 | |
| Nutrients – ASW | 0.000528 | 0.00026 | 68.8 | 2.035 | 0.1116 | |
| Cues – ASW | 0.00138 | 0.000337 | 88.9 | 4.092 | 0.0003 | |
| **Dark Respiration Stationary phase** | | | | | | |
| | Sum Sq | Mean Sq | NumDF | DenDF | F value | Pr (>F) |
| Biov | $9.27 \times 10^{-5}$ | $9.27 \times 10^{-5}$ | 1 | 52 | 14.288 | $4.06 \times 10^{-4}$ (***) |
| Resuspended media | $6.61 \times 10^{-4}$ | $3.30 \times 10^{-4}$ | 2 | 106 | 50.918 | $3.18 \times 10^{-16}$ (***) |
| Post hoc test on resuspended media | | | | | | |
| Resuspended media | EMM | SE | df | Lower CL | Upper CL | |
| Nutrients | 0.01252 | 0.000484 | 106 | 0.01156 | 0.01348 | |
| Cues | 0.00757 | 0.000484 | 106 | 0.00661 | 0.00853 | |
| ASW | 0.00988 | 0.000484 | 106 | 0.00892 | 0.01084 | |
| Contrasts | Estimate | SE | df | t.ratio | p.value | |
| Nutrients – Cues | 0.00494 | 0.00049 | 106 | 10.084 | <.0001 | |
| Nutrients – ASW | 0.00264 | 0.00049 | 106 | 5.378 | <.0001 | |
| Cues – ASW | -0.00231 | 0.00049 | 106 | -4.706 | <.0001 | |

**Table S7.** Linear mixed effect models and post-hoc comparisons testing the relationship between population metabolic rates (μmol $O_2$ $min^{-1}$) and population biovolume ($\mu m^3/\mu l$) in the different media treatments (Resuspended media, 4 levels) for the algal species *Dunaliella tertiolecta* (3 hours exposure) in different growth phases. "Flask_ID" identifying the parent population was included as a random effect in the model. Each metabolic rate (photosynthesis, post-illumination and dark respiration) was analyzed independently. Biov = Biovolume. EMM = estimated marginal means. CL = 95% confidence level. (Biovolume data were divided by $10^5$ before analyses to make the scales of the tested variables more similar).

| **Photosynthesis Exponential phase** | | | | | | |
|---|---|---|---|---|---|---|
| | Sum Sq | Mean Sq | NumDF | DenDF | F value | Pr (>F) |
| Biov | 0.0139435 | 0.0139435 | 1 | 63.975 | 426.6639 | < 2.20 ×10 $^{-16}$ (***) |
| Resuspended media | 0.0005107 | 0.0001702 | 3 | 130.067 | 5.2093 | 0.0020 (**) |
| Biov. × Resuspended media | 0.0006549 | 0.0002183 | 3 | 141.68 | 6.6795 | 0.0003 (***) |
| Post hoc test on biovolume × resuspended media | | | | | | |
| Resuspended media | Slope | SE | df | Lower CL | Upper CL | |
| Nutrients | 0.00570 | 0.000421 | 162 | 0.00487 | 0.00653 | |
| Cues + Nutrients | 0.00548 | 0.000334 | 154 | 0.00482 | 0.00614 | |
| Cues | 0.00492 | 0.000378 | 168 | 0.00417 | 0.00566 | |
| ASW | 0.00384 | 0.000356 | 153 | 0.00314 | 0.00455 | |
| Contrasts | Estimate | SE | df | t.ratio | p.value | |
| Nutrients – (cues + nutrients) | 0.000221 | 0.000474 | 146 | 0.466 | 0.9664 | |
| Nutrients – Cues | 0.00078 | 0.000502 | 144 | 1.552 | 0.4094 | |
| Nutrients – ASW | 0.001856 | 0.00049 | 149 | 3.786 | 0.0013 | |
| (cues + nutrients) – Cues | 0.000559 | 0.000437 | 133 | 1.28 | 0.577 | |
| (cues + nutrients) – ASW | 0.001635 | 0.000423 | 142 | 3.861 | 0.001 | |
| Cues – ASW | 0.001076 | 0.000457 | 142 | 2.353 | 0.0911 | |
| Post hoc test on resuspended media controlling for biovolume (= 3.94 × $10^5$ $\mu m^3/\mu l$) | | | | | | |
| Resuspended media | EMM | SE | df | Lower CL | Upper CL | |
| Nutrients | 0.0351 | 0.00107 | 148 | 0.033 | 0.0372 | |
| Cues + Nutrients | 0.0308 | 0.00102 | 137 | 0.0288 | 0.0328 | |
| Cues | 0.0239 | 0.00105 | 142 | 0.0218 | 0.026 | |
| ASW | 0.0224 | 0.00101 | 138 | 0.0204 | 0.0244 | |
| Contrasts | Estimate | SE | df | t.ratio | p.value | |
| Nutrients – (cues + nutrients) | 0.00433 | 0.00125 | 137 | 3.459 | 0.004 | |
| Nutrients – Cues | 0.01126 | 0.00127 | 137 | 8.844 | <.0001 | |
| Nutrients – ASW | 0.01278 | 0.00125 | 137 | 10.219 | <.0001 | |
| (cues + nutrients) – Cues | 0.00693 | 0.00122 | 127 | 5.696 | <.0001 | |
| (cues + nutrients) - ASW | 0.00845 | 0.00119 | 129 | 7.075 | <.0001 | |
| Cues - ASW | 0.00152 | 0.00122 | 129 | 1.249 | 0.5973 | |
| **Photosynthesis Stationary phase** | | | | | | |
| | Sum Sq | Mean Sq | NumDF | DenDF | F value | Pr (>F) |
| Biov | 0.00023723 | 0.00023723 | 1 | 212.43 | 2.5243 | 0.1136 |
| Resuspended media | 0.00221481 | 0.00073827 | 3 | 187.82 | 7.8558 | 5.76 ×10 $^{-5}$ (***) |
| Post hoc test on resuspended media | | | | | | |
| Resuspended media | EMM | SE | df | Lower CL | Upper CL | |
| Nutrients | 0.0519 | 0.0017 | 150 | 0.0485 | 0.0553 | |
| Cues + Nutrients | 0.0458 | 0.00173 | 155 | 0.0424 | 0.0492 | |
| Cues | 0.0457 | 0.00169 | 148 | 0.0424 | 0.0491 | |
| ASW | 0.0443 | 0.00173 | 155 | 0.0408 | 0.0477 | |

| Contrasts | Estimate | SE | df | t.ratio | | p.value |
|---|---|---|---|---|---|---|
| Nutrients – (cues + nutrients) | 0.006083 | 0.00172 | 189 | 3.529 | | 0.0029 |
| Nutrients – Cues | 0.006192 | 0.00169 | 189 | 3.654 | | 0.0019 |
| Nutrients – ASW | 0.007637 | 0.00172 | 190 | 4.428 | | 0.0001 |
| (cues + nutrients) – Cues | 0.000109 | 0.00171 | 189 | 0.064 | | 0.9999 |
| (cues + nutrients) – ASW | 0.001554 | 0.00174 | 189 | 0.892 | | 0.8091 |
| Cues – ASW | 0.001445 | 0.00172 | 191 | 0.84 | | 0.8354 |
| **Post-Illumination Exponential and Stationary phase** | | | | | | |
| | Sum Sq | Mean Sq | NumDF | DenDF | F value | Pr (>F) |
| $Log_{10}$(Biov) | 0.17698 | 0.17698 | 1 | 88.424 | 1.1957 | 0.27714 |
| Resuspended media | 1.18796 | 0.39599 | 3 | 257.204 | 2.6755 | 0.04771 (*) |
| $Log_{10}$(Biov) × Resuspended media | 1.01567 | 0.33856 | 3 | 257.675 | 2.2874 | 0.07905 (.) |
| Post hoc test on resuspended media controlling for biovolume (= $1×10^{6}$ $\mu m^3/\mu l$) | | | | | | |
| Resuspended media | EMM | SE | df | Lower CL | | Upper CL |
| Nutrients | -1.47 | 0.0403 | 346 | -1.55 | | -1.39 |
| Cues + nutrients | -1.88 | 0.041 | 347 | -1.96 | | -1.8 |
| Cues | -2.41 | 0.0521 | 348 | -2.51 | | -2.31 |
| ASW | -2.22 | 0.0456 | 347 | -2.31 | | -2.14 |
| Contrasts | Estimate | SE | df | t.ratio | | p.value |
| Nutrients – (cues + nutrients) | 0.415 | 0.0562 | 252 | 7.377 | | <.0001 |
| Nutrients – Cues | 0.939 | 0.0648 | 282 | 14.494 | | <.0001 |
| Nutrients – ASW | 0.755 | 0.0597 | 267 | 12.652 | | <.0001 |
| (Cues + nutrients) – cues | 0.525 | 0.0652 | 284 | 8.046 | | <.0001 |
| (Cues + nutrients) – ASW | 0.341 | 0.0601 | 267 | 5.667 | | <.0001 |
| Cues – ASW | -0.184 | 0.0681 | 279 | -2.704 | | 0.0363 |
| **Dark Respiration Exponential phase** | | | | | | |
| | Sum Sq | Mean Sq | NumDF | DenDF | F value | Pr (>F) |
| Biov | $2.23×10^{-5}$ | $2.23×10^{-5}$ | 1 | 56.578 | 4.201 | 0.0450 (*) |
| Resuspended media | $6.56×10^{-5}$ | $2.19×10^{-5}$ | 3 | 97.957 | 4.1254 | 0.0084 (**) |
| Post hoc test on resuspended media | | | | | | |
| Resuspended media | EMM | SE | df | Lower CL | | Upper CL |
| Nutrients | 0.00428 | 0.0005 | 124 | 0.00329 | | 0.00527 |
| Cues + nutrients | 0.00448 | 0.000568 | 133 | 0.00336 | | 0.0056 |
| Cues | 0.00443 | 0.000564 | 134 | 0.00331 | | 0.00554 |
| ASW | 0.00281 | 0.000463 | 108 | 0.0019 | | 0.00373 |
| Contrasts | Estimate | SE | df | t.ratio | | p.value |
| Nutrients – (Cues + nutrients) | $-1.96×10^{-4}$ | 0.000635 | 102.7 | -0.308 | | 0.9898 |
| Nutrients – Cues | $-1.41×10^{-4}$ | 0.000623 | 98.3 | -0.227 | | 0.9959 |
| Nutrients – ASW | $1.47×10^{-3}$ | 0.00055 | 99.7 | 2.672 | | 0.043 |
| (Cues + nutrients) – Cues | $5.45×10^{-5}$ | 0.000656 | 93.8 | 0.083 | | 0.9998 |
| (Cues + nutrients) - ASW | $1.67×10^{-3}$ | 0.00061 | 104.6 | 2.73 | | 0.0368 |
| Cues - ASW | $1.61×10^{-3}$ | 0.000605 | 100.7 | 2.66 | | 0.0443 |
| **Dark Respiration Stationary phase** | | | | | | |
| | Sum Sq | Mean Sq | NumDF | DenDF | F value | Pr (>F) |
| Biov | 0.0003108 | 0.00031077 | 1 | 136.4 | 10.622 | 0.0014 (**) |
| Resuspended media | 0.0047732 | 0.00159105 | 3 | 168.13 | 54.38 | $<2.2×10^{-16}$ (***) |
| Post hoc test on resuspended media | | | | | | |
| Resuspended media | EMM | SE | df | Lower CL | | Upper CL |

| | | | | | |
|---|---|---|---|---|---|
| Nutrients | 0.0206 | 0.00085 | 200 | 0.01896 | 0.0223 |
| Cues + nutrients | 0.0151 | 0.000885 | 207 | 0.01332 | 0.0168 |
| Cues | 0.0085 | 0.000797 | 187 | 0.00692 | 0.0101 |
| ASW | 0.0116 | 0.00082 | 192 | 0.01001 | 0.0132 |
| Contrasts | Estimate | SE | df | t.ratio | p.value |
| Nutrients – (Cues + nutrients) | 0.00557 | 0.00105 | 173 | 5.31 | <.0001 |
| Nutrients – Cues | 0.01214 | 0.000989 | 173 | 12.279 | <.0001 |
| Nutrients – ASW | 0.00901 | 0.001 | 175 | 8.962 | <.0001 |
| (Cues + nutrients) – Cues | 0.00657 | 0.00102 | 174 | 6.456 | <.0001 |
| (Cues + nutrients) – ASW | 0.00343 | 0.00061 | 176 | 3.321 | 0.0059 |
| Cues – ASW | -0.00313 | 0.000964 | 172 | -3.252 | 0.0075 |

**Table S8.** Linear mixed effect models and post-hoc comparisons testing the relationship between population metabolic rates (µmol $O_2$ $min^{-1}$) and population biovolume ($\mu m^3/\mu l$) in the different media treatments (Resuspended media, 4 levels) for the algal species *Tetraselmis sp.* (3 hours exposure). "Flask_ID" identifying the parent population was included as a random effect in the model. Each metabolic rate (photosynthesis, post-illumination and dark respirations) was analyzed independently. Biov = Biovolume. EMM = estimated marginal means. CL = 95% confidence level. (Biovolume data were divided by $10^5$ before analyses to make the scales of the tested variables more similar).

| **Photosynthesis Exponential phase** | | | | | | |
|---|---|---|---|---|---|---|
| | Sum Sq | Mean Sq | NumDF | DenDF | F value | Pr (>F) |
| $Log_{10}$(Biov) | 0.76811 | 0.76811 | 1 | 128.93 | 39.6301 | $4.43\times10^{-9}$ (***) |
| Resuspended media | 0.16586 | 0.05529 | 3 | 203.02 | 2.8525 | 0.0384 (*) |
| $Log_{10}$(Biov) × Resuspended media | 0.15616 | 0.05205 | 3 | 202.99 | 2.6857 | 0.0477 (*) |
| Post hoc test on biovolume × resuspended media | | | | | | |
| Resuspended media | Slope | SE | df | Lower CL | Upper CL | |
| Nutrients | 0.386 | 0.091 | 202 | 0.206 | 0.566 | |
| Cues + Nutrients | 0.558 | 0.095 | 209 | 0.370 | 0.746 | |
| Cues | 0.548 | 0.095 | 205 | 0.361 | 0.735 | |
| ASW | 0.582 | 0.104 | 199 | 0.377 | 0.788 | |
| Contrasts | Estimate | SE | df | t.ratio | p.value | |
| Nutrients – (cues + nutrients) | -0.1719 | 0.0773 | 206 | -2.223 | 0.1205 | |
| Nutrients – Cues | -0.1619 | 0.0768 | 206 | -2.107 | 0.1543 | |
| Nutrients – ASW | -0.1963 | 0.0811 | 213 | -2.42 | 0.0765 | |
| (cues + nutrients) – Cues | 0.0100 | 0.0794 | 206 | 0.126 | 0.9993 | |
| (cues + nutrients) – ASW | -0.0244 | 0.0839 | 211 | -0.291 | 0.9914 | |
| Cues – ASW | -0.0344 | 0.0824 | 210 | -0.418 | 0.9754 | |
| Post hoc test on resuspended media controlling for biovolume (= $4.54\times10^5$ $\mu m^3/\mu l$) | | | | | | |
| Resuspended media | EMM | SE | df | Lower CL | Upper CL | |
| Nutrients | -0.998 | 0.0333 | 115 | -1.06 | -0.932 | |
| Cues + Nutrients | -0.949 | 0.0336 | 120 | -1.02 | -0.882 | |
| Cues | -1.06 | 0.0329 | 114 | -1.13 | -0.995 | |
| ASW | -1.027 | 0.0327 | 111 | -1.09 | -0.962 | |
| Contrasts | Estimate | SE | df | t.ratio | p.value | |
| Nutrients – (cues + nutrients) | -0.0493 | 0.025 | 205 | -1.977 | 0.2001 | |
| Nutrients – Cues | 0.0617 | 0.0245 | 208 | 2.516 | 0.0604 | |
| Nutrients – ASW | 0.0284 | 0.0243 | 207 | 1.168 | 0.6479 | |
| (cues + nutrients) – Cues | 0.111 | 0.0249 | 207 | 4.451 | 0.0001 | |
| (cues + nutrients) – ASW | 0.0777 | 0.0249 | 207 | 3.124 | 0.0109 | |
| Cues – ASW | -0.0333 | 0.0241 | 206 | -1.383 | 0.5113 | |
| **Post-Illumination Exponential phase** | | | | | | |
| | Sum Sq | Mean Sq | NumDF | DenDF | F value | Pr (>F) |
| $Log_{10}$(Biov) | 2.94375 | 2.94375 | 1 | 95.683 | 152.9433 | $<2.2\times10^{-16}$ (***) |
| Resuspended media | 0.24153 | 0.08051 | 3 | 206.38 | 4.1829 | 0.0067 (**) |
| $Log_{10}$(Biov) × Resuspended media | 0.20257 | 0.06752 | 3 | 206.494 | 3.5081 | 0.0163 (*) |
| Post hoc test on biovolume × resuspended media | | | | | | |
| Resuspended media | Slope | SE | df | Lower CL | Upper CL | |
| Nutrients | 0.613 | 0.0723 | 208 | 0.471 | 0.756 | |
| Cues + nutrients | 0.701 | 0.0760 | 218 | 0.551 | 0.851 | |
| Cues | 0.723 | 0.0752 | 212 | 0.574 | 0.871 | |
| ASW | 0.872 | 0.0823 | 205 | 0.709 | 1.034 | |
| Contrasts | Estimate | SE | df | t.ratio | p.value | |

| | | | | | | |
|---|---|---|---|---|---|---|
| Nutrients – (cues + nutrients) | -0.0876 | 0.0770 | 207 | -1.138 | | 0.6666 |
| Nutrients – Cues | -0.1094 | 0.0764 | 208 | -1.433 | | 0.4805 |
| Nutrients – ASW | -0.2587 | 0.0802 | 212 | -3.226 | | 0.0079 |
| (Cues + nutrients) – cues | -0.0219 | 0.0790 | 208 | -0.277 | | 0.9926 |
| (Cues + nutrients) – ASW | -0.1712 | 0.0830 | 212 | -2.063 | | 0.1687 |
| Cues – ASW | -0.1493 | 0.0816 | 210 | -1.829 | | 0.2627 |
| Post hoc test on resuspended media controlling for biovolume (= 4.54 × $10^5$ $\mu m^3/\mu l$) | | | | | | |
| Resuspended media | EMM | SE | df | Lower CL | | Upper CL |
| Nutrients | -1.42 | 0.0245 | 165 | -1.47 | | -1.37 |
| Cues + nutrients | -1.4 | 0.025 | 174 | -1.45 | | -1.35 |
| Cues | -1.57 | 0.0242 | 166 | -1.61 | | -1.52 |
| ASW | -1.51 | 0.0239 | 160 | -1.56 | | -1.47 |
| Contrasts | Estimate | SE | df | t.ratio | | p.value |
| Nutrients – (Cues + nutrients) | -0.0194 | 0.0248 | 207 | -0.782 | | 0.8628 |
| Nutrients – Cues | 0.1477 | 0.0244 | 208 | 6.062 | | <.0001 |
| Nutrients – ASW | 0.0956 | 0.0242 | 209 | 3.955 | | 0.0006 |
| (Cues + nutrients) – Cues | 0.1671 | 0.0248 | 208 | 6.739 | | <.0001 |
| (Cues + nutrients) – ASW | 0.115 | 0.0247 | 210 | 4.654 | | <.0001 |
| Cues – ASW | -0.0521 | 0.024 | 207 | -2.177 | | 0.1332 |
| **Dark Respiration Exponential phase** | | | | | | |
| | Sum Sq | Mean Sq | NumDF | DenDF | F value | Pr (>F) |
| Biov | $1.22 \times10^{-4}$ | $1.22\times10^{-4}$ | 1 | 103.61 | 52.8534 | $7.04\times10^{-11}$ (***) |
| Resuspended media | $2.27 \times10^{-5}$ | $7.56\times10^{-6}$ | 3 | 171.81 | 3.276 | 0.0224 (*) |
| Biov × Resuspended media | $2.18 \times10^{-5}$ | $7.28\times10^{-6}$ | 3 | 173.43 | 3.1549 | 0.0262 (*) |
| Post hoc test on biovolume × resuspended media | | | | | | |
| Resuspended media | Slope | SE | df | Lower CL | | Upper CL |
| Nutrients | 0.000495 | $9.20\times10^{-5}$ | 207 | $3.13\times10^{-4}$ | | 0.000676 |
| Cues + nutrients | 0.000566 | $1.24\times10^{-4}$ | 216 | $3.21\times10^{-4}$ | | 0.000811 |
| Cues | 0.000204 | $1.08\times10^{-4}$ | 215 | $-8.21\times10^{-6}$ | | 0.000417 |
| ASW | 0.000569 | $8.83\times10^{-5}$ | 202 | $3.95\times10^{-4}$ | | 0.000743 |
| Contrasts | Estimate | SE | df | t.ratio | | p.value |
| Nutrients – (Cues + nutrients) | $-7.13\times10^{-5}$ | 0.000142 | 179 | -0.501 | | 0.9588 |
| Nutrients – Cues | $2.90\times10^{-4}$ | 0.000132 | 184 | 2.202 | | 0.1265 |
| Nutrients – ASW | $-7.45\times10^{-5}$ | 0.000114 | 176 | -0.653 | | 0.9144 |
| (Cues + nutrients) – Cues | $3.62\times10^{-4}$ | 0.000152 | 173 | 2.382 | | 0.0843 |
| (Cues + nutrients) - ASW | $-3.17\times10^{-6}$ | 0.000138 | 168 | -0.023 | | 1 |
| Cues – ASW | $-3.65\times10^{-4}$ | 0.000127 | 171 | -2.873 | | 0.0235 |
| Post hoc test on resuspended media controlling for biovolume (= 4.85 × $10^5$ $\mu m^3/\mu l$) | | | | | | |
| Resuspended media | EMM | SE | df | Lower CL | | Upper CL |
| Nutrients | 0.00332 | 0.000227 | 195 | 0.00288 | | 0.00377 |
| Cues + nutrients | 0.00368 | 0.000262 | 210 | 0.00316 | | 0.0042 |
| Cues | 0.00359 | 0.000266 | 211 | 0.00307 | | 0.00412 |
| ASW | 0.00308 | 0.000217 | 187 | 0.00265 | | 0.00351 |
| Contrasts | Estimate | SE | df | t.ratio | | p.value |
| Nutrients – (Cues + nutrients) | $-3.56\times10^{-4}$ | 0.000307 | 163 | -1.158 | | 0.6543 |
| Nutrients – Cues | $-2.70\times10^{-4}$ | 0.000311 | 164 | -0.87 | | 0.8206 |
| Nutrients – ASW | $2.41\times10^{-4}$ | 0.000273 | 159 | 0.884 | | 0.8132 |
| (Cues + nutrients) – Cues | $8.56\times10^{-5}$ | 0.000331 | 158 | 0.259 | | 0.9939 |
| (Cues + nutrients) – ASW | $5.97\times10^{-4}$ | 0.000299 | 159 | 1.995 | | 0.1943 |
| Cues – ASW | $5.11\times10^{-4}$ | 0.000302 | 160 | 1.691 | | 0.3319 |

**Table S9.** *Nannochloropsis granulata* (3 hours exposure). Linear mixed effect models and post-hoc comparisons testing the relationship between population metabolic rates (µmol $O_2$ $min^{-1}$) and population biovolume ($µm^3$/µl) in the different media treatments (Resuspended media, 4 levels) in different growth phases. "Flask_ID" identifying the parent population was included as a random effect in the model. Each metabolic rate (photosynthesis, post-illumination and dark respirations) was analyzed independently. Biov = Biovolume. EMM = estimated marginal means. CL = 95% confidence level. (Biovolume data were divided by $10^5$ before analyses to make the scales of the tested variables more similar).

| **Photosynthesis Exponential phase** | | | | | | |
|---|---|---|---|---|---|---|
| | Sum Sq | Mean Sq | NumDF | DenDF | F value | Pr (>F) |
| Biov | 0.0141663 | 0.0141663 | 1 | 95.103 | 673.3663 | $< 2 \times 10^{-16}$ (***) |
| Resuspended media | 0.0000209 | 0.000007 | 3 | 241.09 | 5.2093 | 0.8034 |
| Biov × Resuspended media | 0.0002397 | 0.0000799 | 3 | 246.597 | 6.6795 | 0.0109 (*) |
| Post hoc test on biovolume × resuspended media | | | | | | |
| Resuspended media | Slope | SE | df | Lower CL | Upper CL | |
| Nutrients | $4.7 \times 10^{-3}$ | $2.44 \times 10^{-4}$ | 219 | $4.22\times10^{-3}$ | $5.18\times10^{-3}$ | |
| Cues + Nutrients | $5.2 \times 10^{-3}$ | $2.42\times10^{-4}$ | 210 | 0.00472 | 0.00567 | |
| Cues | $4.9 \times 10^{-3}$ | $2.51\times10^{-4}$ | 239 | $4.38\times10^{-3}$ | $5.37\times10^{-3}$ | |
| ASW | $5.5\times10^{-3}$ | $2.55\times10^{-4}$ | 232 | $4.97\times10^{-3}$ | $5.97\times10^{-3}$ | |
| Contrasts | Estimate | SE | df | t.ratio | p.value | |
| Nutrients – (cues + nutrients) | -0.000497 | 0.000242 | 259 | -2.055 | 0.1709 | |
| Nutrients – Cues | $-1.75\times10^{-4}$ | $2.50\times10^{-4}$ | 255 | -0.702 | 0.8963 | |
| Nutrients – ASW | $-7.73\times10^{-4}$ | $2.50\times10^{-4}$ | 255 | -3.092 | 0.0118 | |
| (cues + nutrients) – Cues | 0.000322 | 0.00025 | 260 | 1.288 | 0.5713 | |
| (cues + nutrients) – ASW | -0.000276 | 0.00025 | 259 | -1.105 | 0.6867 | |
| Cues – ASW | $5.98\times10^{-4}$ | $2.53\times10^{-4}$ | 250 | -2.36 | 0.0877 | |
| Post hoc test on resuspended media controlling for biovolume (= $4.61 \times 10^5$ $µm^3$/µl) | | | | | | |
| Resuspended media | EMM | SE | df | Lower CL | Upper CL | |
| Nutrients | 0.0257 | 0.000727 | 187 | 0.0242 | 0.0271 | |
| Cues + Nutrients | 0.0273 | 0.000722 | 183 | 0.0259 | 0.0287 | |
| Cues | 0.0252 | 0.000737 | 190 | 0.0237 | 0.0266 | |
| ASW | 0.0284 | 0.000734 | 190 | 0.0269 | 0.0298 | |
| Contrasts | Estimate | SE | df | t.ratio | p.value | |
| Nutrients – (cues + nutrients) | -0.001628 | 0.000696 | 252 | -2.338 | 0.0923 | |
| Nutrients – Cues | 0.000492 | 0.000707 | 252 | 0.697 | 0.8983 | |
| Nutrients – ASW | -0.002702 | 0.000704 | 251 | -3.841 | 0.0009 | |
| (cues + nutrients) – Cues | 0.00212 | 0.000704 | 253 | 3.012 | 0.0151 | |
| (cues + nutrients) – ASW | -0.001074 | 0.000702 | 252 | -1.531 | 0.4206 | |
| Cues - ASW | -0.003195 | 0.000707 | 250 | -4.518 | 0.0001 | |
| **Photosynthesis Stationary phase** | | | | | | |
| | Sum Sq | Mean Sq | NumDF | DenDF | F value | Pr (>F) |
| Biov | 0.00004524 | $4.52\times10^{-5}$ | 1 | 60.203 | 0.3133 | 0.5777 |
| Resuspended media | 0.00068597 | $2.29\times10^{-4}$ | 3 | 99.116 | 1.5838 | 0.1981 |
| **Post-Illumination Exponential phase** | | | | | | |
| | Sum Sq | Mean Sq | NumDF | DenDF | F value | Pr (>F) |
| Biov | 0.00052379 | 0.00052379 | 1 | 93.452 | 92.8938 | $1.14\times10^{-15}$ (***) |
| Resuspended media | $3.446\times10^{-5}$ | $1.149\times10^{-5}$ | 3 | 240.88 | 2.0372 | 0.1093 |
| **Post-Illumination Stationary phase** | | | | | | |
| | Sum Sq | Mean Sq | NumDF | DenDF | F value | Pr (>F) |
| Biov | 0.00008952 | $8.95 \times10^{-5}$ | 1 | 71.586 | 1.8028 | 0.18362 |
| Resuspended media | 0.00043424 | $1.45 \times10^{-4}$ | 3 | 94.751 | 2.915 | 0.0383 (*) |
| Post hoc test on resuspended media | | | | | | |

| Resuspended media | EMM | SE | df | Lower CL | Upper CL | |
|---|---|---|---|---|---|---|
| Nutrients | 0.0161 | 0.00166 | 85.8 | 0.01285 | 0.0194 | |
| Cues + nutrients | 0.0165 | 0.00169 | 91 | 0.01311 | 0.0198 | |
| Cues | 0.0123 | 0.00165 | 86.1 | 0.00901 | 0.0156 | |
| ASW | 0.017 | 0.00163 | 82.2 | 0.01372 | 0.0202 | |
| Contrasts | Estimate | SE | df | t.ratio | p.value | |
| Nutrients – (Cues + nutrients) | -0.000325 | 0.00178 | 93.3 | -0.182 | 0.9978 | |
| Nutrients – Cues | 0.003844 | 0.00178 | 98.2 | 2.163 | 0.141 | |
| Nutrients – ASW | -0.000832 | 0.00173 | 93.3 | -0.48 | 0.9633 | |
| (Cues + nutrients) – Cues | 0.00417 | 0.00181 | 95.8 | 2.31 | 0.1029 | |
| (Cues + nutrients) - ASW | -0.000506 | 0.00179 | 95 | -0.283 | 0.992 | |
| Cues - ASW | -0.004676 | 0.00175 | 99.3 | -2.667 | 0.0436 | |
| **Dark Respiration Exponential phase** | | | | | | |
| | Sum Sq | Mean Sq | NumDF | DenDF | F value | Pr (>F) |
| Biov | $9.08\times10^{-5}$ | $9.08\times10^{-5}$ | 1 | 102.9 | 67.664 | $6.27\times10^{-13}$ (***) |
| Resuspended media | $7.55\times10^{-5}$ | $2.52\times10^{-5}$ | 3 | 235 | 18.746 | $6.16\times10^{-11}$ (***) |
| Post hoc test on resuspended media | | | | | | |
| Resuspended media | EMM | SE | df | Lower CL | Upper CL | |
| Nutrients | 0.00404 | 0.0002 | 163 | 0.00365 | 0.00444 | |
| Cues + nutrients | 0.00411 | 0.000204 | 172 | 0.00371 | 0.00451 | |
| Cues | 0.00371 | 0.000208 | 180 | 0.0033 | 0.00412 | |
| ASW | 0.00291 | 0.000201 | 166 | 0.00251 | 0.0033 | |
| Contrasts | Estimate | SE | df | t.ratio | p.value | |
| Nutrients – (Cues + nutrients) | $-6.53\times10^{-5}$ | 0.000182 | 240 | -0.359 | 0.9841 | |
| Nutrients – Cues | $3.30\times10^{-4}$ | 0.000185 | 241 | 1.786 | 0.2828 | |
| Nutrients – ASW | $1.14\times10^{-3}$ | 0.000178 | 237 | 6.391 | <.0001 | |
| (Cues + nutrients) – Cues | $3.96\times10^{-4}$ | 0.000188 | 241 | 2.1 | 0.1562 | |
| (Cues + nutrients) – ASW | $1.20\times10^{-3}$ | 0.000183 | 240 | 6.552 | <.0001 | |
| Cues – ASW | $8.05\times10^{-4}$ | 0.000185 | 238 | 4.352 | 0.0001 | |
| **Dark Respiration Stationary phase** | | | | | | |
| | Sum Sq | Mean Sq | NumDF | DenDF | F value | Pr (>F) |
| Biov | $1.07\times10^{-5}$ | $1.07\times10^{-5}$ | 1 | 91.174 | 1.4178 | 0.2368 |
| Resuspended media | $1.77\times10^{-4}$ | $5.90\times10^{-5}$ | 3 | 93.036 | 7.8446 | 0.0001 (***) |
| Post hoc test on resuspended media | | | | | | |
| Resuspended media | EMM | SE | df | Lower CL | Upper CL | |
| Nutrients | 0.0081 | 0.000772 | 65.9 | 0.00656 | 0.00964 | |
| Cues + nutrients | 0.00609 | 0.000796 | 73 | 0.0045 | 0.00767 | |
| Cues | 0.00482 | 0.000764 | 64.6 | 0.0033 | 0.00635 | |
| ASW | 0.00686 | 0.000767 | 63.7 | 0.00533 | 0.00839 | |
| Contrasts | Estimate | SE | df | t.ratio | p.value | |
| Nutrients – (Cues + nutrients) | 0.002016 | 0.000712 | 92.1 | 2.832 | 0.0286 | |
| Nutrients – Cues | 0.003278 | 0.000693 | 96.8 | 4.732 | <.0001 | |
| Nutrients – ASW | 0.001245 | 0.000675 | 91.5 | 1.845 | 0.2594 | |
| (Cues + nutrients) – Cues | 0.001262 | 0.000716 | 94 | 1.763 | 0.2977 | |
| (Cues + nutrients) – ASW | -0.000771 | 0.000715 | 93.5 | -1.079 | 0.7034 | |
| Cues – ASW | -0.002033 | 0.000685 | 98.4 | -2.966 | 0.0195 | |

**Table S10.** Linear mixed effect models and post-hoc comparisons testing the relationship between average cell volume ($\mu m^3$) and experiment day in the different media treatments (Resuspended media, 4 levels) for the algal species *Tetraselmis sp.* (3 hours exposure). "Flask_ID" identifying the parent population was included as a random effect in the model. Exp Day = Experiment Day. Vol = Average cell volume. EMM = estimated marginal means. CL = 95% confidence level.

| ***Tetraselmis* Average cell volume** | | | | | | |
|---|---|---|---|---|---|---|
| | Sum Sq | Mean Sq | NumDF | DenDF | F value | Pr (>F) |
| Exp Day | 240508 | 34358 | 7 | 72 | 86.110 | $<2.2 \times 10^{-16}$ (***) |
| Resuspended media | 22946 | 7649 | 3 | 237 | 19.169 | $3.656 \times 10^{-11}$ (***) |
| Post hoc test on resuspended media interaction | | | | | | |
| Resuspended media | EMM | SE | df | Lower CL | Upper CL | |
| Nutrients | 656 | 4.21 | 113 | 647 | 664 | |
| Cues + Nutrients | 665 | 4.21 | 113 | 657 | 674 | |
| Cues | 678 | 4.21 | 113 | 670 | 687 | |
| ASW | 673 | 4.21 | 113 | 665 | 682 | |
| Contrasts | Estimate | SE | df | t.ratio | p.value | |
| Nutrients – (cues + nutrients) | -9.53 | 3.16 | 237 | -3.018 | 0.0149 | |
| Nutrients – Cues | -22.40 | 3.16 | 237 | -7.092 | <0.0001 | |
| Nutrients – ASW | -17.39 | 3.16 | 237 | -5.505 | <0.0001 | |
| (Cues + nutrients) – Cues | -12.87 | 3.16 | 237 | -4.074 | 0.0004 | |
| (Cues + nutrients) – ASW | -7.85 | 3.16 | 237 | -2.486 | 0.0646 | |
| Cues – ASW | 5.01 | 3.16 | 237 | 1.588 | 0.3876 | |

**Table S11.** Linear mixed effect models and post-hoc comparisons testing the relationship between average cell volume (µm$^3$) and experiment day in the different media treatments (Resuspended media, 4 levels) for the algal species *Dunaliella tertiolecta* (3 hours exposure). "Flask_ID" identifying the parent population was included as a random effect in the model. Exp Day = Experiment Day. Vol = Average cell volume. EMM = estimated marginal means. CL = 95% confidence level.

| ***Dunaliella* Average cell volume** | | | | | | |
|---|---|---|---|---|---|---|
| | Sum Sq | Mean Sq | NumDF | DenDF | F value | Pr (>F) |
| Exp Day | 77161 | 7716.1 | 10 | 99 | 52.1739 | <2.2 $\times 10^{-16}$ (***) |
| Resuspended media | 33853 | 11284.4 | 3 | 297 | 76.3016 | <2.2 $\times 10^{-16}$ (***) |
| Exp Day × Resuspended media | 20602 | 686.7 | 30 | 297 | 4.6434 | 1.289 $\times 10^{-12}$ (***) |
| Post hoc test on experiment day × resuspended media | | | | | | |
| **Experiment Day = 1** | | | | | | |
| Resuspended media | EMM | SE | df | Lower CL | Upper CL | |
| Nutrients | 454 | 8.12 | 141 | 438 | 470 | |
| Cues + Nutrients | 454 | 8.12 | 141 | 438 | 470 | |
| Cues | 472 | 8.12 | 141 | 456 | 488 | |
| ASW | 448 | 8.12 | 141 | 432 | 464 | |
| Contrasts | Estimate | SE | df | t.ratio | p.value | |
| Nutrients – (cues + nutrients) | -0.322 | 5.44 | 297 | -0.059 | 0.9999 | |
| Nutrients – Cues | -17.882 | 5.44 | 297 | -3.288 | 0.0062 | |
| Nutrients – ASW | 5.701 | 5.44 | 297 | 1.048 | 0.7212 | |
| (cues + nutrients) – Cues | -17.561 | 5.44 | 297 | -3.229 | 0.0075 | |
| (cues + nutrients) – ASW | 6.023 | 5.44 | 297 | 1.107 | 0.6853 | |
| Cues – ASW | 23.583 | 5.44 | 297 | 4.336 | 0.0001 | |
| **Experiment Day = 2** | | | | | | |
| Resuspended media | EMM | SE | df | Lower CL | Upper CL | |
| Nutrients | 425 | 8.12 | 141 | 409 | 441 | |
| Cues + Nutrients | 449 | 8.12 | 141 | 433 | 465 | |
| Cues | 461 | 8.12 | 141 | 445 | 477 | |
| ASW | 458 | 8.12 | 141 | 442 | 474 | |
| Contrasts | Estimate | SE | df | t.ratio | p.value | |
| Nutrients – (cues + nutrients) | -23.788 | 5.44 | 297 | -4.374 | 0.0001 | |
| Nutrients – Cues | -35.921 | 5.44 | 297 | -6.605 | <0.0001 | |
| Nutrients – ASW | -32.671 | 5.44 | 297 | -6.007 | <0.0001 | |
| (cues + nutrients) – Cues | -12.133 | 5.44 | 297 | -2.231 | 0.1173 | |
| (cues + nutrients) – ASW | -8.883 | 5.44 | 297 | -1.633 | 0.3615 | |
| Cues - ASW | 3.251 | 5.44 | 297 | 0.598 | 0.9327 | |
| **Experiment Day = 3** | | | | | | |
| Resuspended media | EMM | SE | df | Lower CL | Upper CL | |
| Nutrients | 406 | 8.12 | 141 | 390 | 422 | |
| Cues + Nutrients | 452 | 8.12 | 141 | 436 | 468 | |
| Cues | 465 | 8.12 | 141 | 449 | 481 | |
| ASW | 445 | 8.12 | 141 | 428 | 461 | |
| Contrasts | Estimate | SE | df | t.ratio | p.value | |
| Nutrients – (cues + nutrients) | -45.690 | 5.44 | 297 | -8.401 | <0.0001 | |
| Nutrients – Cues | -58.868 | 5.44 | 297 | -10.824 | <0.0001 | |
| Nutrients – ASW | -38.268 | 5.44 | 297 | -7.036 | <0.0001 | |
| (cues + nutrients) – Cues | -13.178 | 5.44 | 297 | -2.423 | 0.0750 | |
| (cues + nutrients) – ASW | 7.422 | 5.44 | 297 | 1.365 | 0.5225 | |
| Cues - ASW | 20.600 | 5.44 | 297 | 3.788 | 0.0011 | |
| **Experiment Day = 4** | | | | | | |
| Resuspended media | EMM | SE | df | Lower CL | Upper CL | |
| Nutrients | 386 | 8.12 | 141 | 370 | 402 | |

| | | | | | |
|---|---|---|---|---|---|
| Cues + Nutrients | 415 | 8.12 | 141 | 399 | 431 |
| Cues | 425 | 8.12 | 141 | 409 | 441 |
| ASW | 416 | 8.12 | 141 | 400 | 432 |
| Contrasts | Estimate | SE | df | t.ratio | p.value |
| Nutrients – (cues + nutrients) | -29.157 | 5.44 | 297 | -5.361 | <0.0001 |
| Nutrients – Cues | -38.369 | 5.44 | 297 | -7.055 | <0.0001 |
| Nutrients – ASW | -29.506 | 5.44 | 297 | -5.425 | <0.0001 |
| (cues + nutrients) – Cues | -9.212 | 5.44 | 297 | -1.694 | 0.3288 |
| (cues + nutrients) – ASW | -0.349 | 5.44 | 297 | -0.064 | 0.9999 |
| Cues - ASW | 8.863 | 5.44 | 297 | 1.630 | 0.3635 |
| **Experiment Day = 6** | | | | | |
| Resuspended media | EMM | SE | df | Lower CL | Upper CL |
| Nutrients | 394 | 8.12 | 141 | 378 | 410 |
| Cues + Nutrients | 411 | 8.12 | 141 | 395 | 427 |
| Cues | 429 | 8.12 | 141 | 413 | 445 |
| ASW | 413 | 8.12 | 141 | 397 | 429 |
| Contrasts | Estimate | SE | df | t.ratio | p.value |
| Nutrients – (cues + nutrients) | -16.918 | 5.44 | 297 | -3.111 | 0.0110 |
| Nutrients – Cues | -35.356 | 5.44 | 297 | -6.501 | <0.0001 |
| Nutrients – ASW | -18.785 | 5.44 | 297 | -3.454 | 0.0035 |
| (cues + nutrients) – Cues | -18.438 | 5.44 | 297 | -3.390 | 0.0044 |
| (cues + nutrients) – ASW | -1.867 | 5.44 | 297 | -0.343 | 0.9861 |
| Cues - ASW | 16.571 | 5.44 | 297 | 3.047 | 0.0134 |
| **Experiment Day = 7** | | | | | |
| Resuspended media | EMM | SE | df | Lower CL | Upper CL |
| Nutrients | 451 | 8.12 | 141 | 434 | 467 |
| Cues + Nutrients | 464 | 8.12 | 141 | 448 | 480 |
| Cues | 469 | 8.12 | 141 | 453 | 485 |
| ASW | 466 | 8.12 | 141 | 450 | 482 |
| Contrasts | Estimate | SE | df | t.ratio | p.value |
| Nutrients – (cues + nutrients) | -13.432 | 5.44 | 297 | -2.470 | 0.0668 |
| Nutrients – Cues | -18.369 | 5.44 | 297 | -3.377 | 0.0046 |
| Nutrients – ASW | -15.042 | 5.44 | 297 | -2.766 | 0.0306 |
| (cues + nutrients) – Cues | -4.936 | 5.44 | 297 | -0.908 | 0.8008 |
| (cues + nutrients) – ASW | -1.610 | 5.44 | 297 | -0.296 | 0.9910 |
| Cues - ASW | 3.326 | 5.44 | 297 | 0.612 | 0.9238 |
| **Experiment Day = 9** | | | | | |
| Resuspended media | EMM | SE | df | Lower CL | Upper CL |
| Nutrients | 531 | 8.12 | 141 | 515 | 547 |
| Cues + Nutrients | 538 | 8.12 | 141 | 522 | 554 |
| Cues | 531 | 8.12 | 141 | 515 | 547 |
| ASW | 546 | 8.12 | 141 | 530 | 562 |
| Contrasts | Estimate | SE | df | t.ratio | p.value |
| Nutrients – (cues + nutrients) | -7.172 | 5.44 | 297 | -1.319 | 0.5517 |
| Nutrients – Cues | -0.236 | 5.44 | 297 | -0.043 | 1.0000 |
| Nutrients – ASW | -14.943 | 5.44 | 297 | -2.748 | 0.0322 |
| (cues + nutrients) – Cues | 6.936 | 5.44 | 297 | 1.275 | 0.5794 |
| (cues + nutrients) – ASW | -7.770 | 5.44 | 297 | -1.429 | 0.4824 |
| Cues - ASW | -14.706 | 5.44 | 297 | -2.704 | 0.0363 |
| **Experiment Day = 10** | | | | | |
| Resuspended media | EMM | SE | df | Lower CL | Upper CL |
| Nutrients | 494 | 8.12 | 141 | 478 | 511 |
| Cues + Nutrients | 512 | 8.12 | 141 | 496 | 528 |
| Cues | 510 | 8.12 | 141 | 494 | 526 |
| ASW | 517 | 8.12 | 141 | 501 | 533 |
| Contrasts | Estimate | SE | df | t.ratio | p.value |
| Nutrients – (cues + nutrients) | -17.515 | 5.44 | 297 | -3.220 | 0.0077 |
| Nutrients – Cues | -15.130 | 5.44 | 297 | -2.782 | 0.0292 |
| Nutrients – ASW | -22.184 | 5.44 | 297 | -4.079 | 0.0003 |

| | | | | | |
|---|---|---|---|---|---|
| (cues + nutrients) – Cues | 2.385 | 5.44 | 297 | 0.439 | 0.9717 |
| (cues + nutrients) – ASW | -4.669 | 5.44 | 297 | -0.859 | 0.8261 |
| Cues - ASW | -7.055 | 5.44 | 297 | -1.297 | 0.5655 |
| **Experiment Day = 11** | | | | | |
| Resuspended media | EMM | SE | df | Lower CL | Upper CL |
| Nutrients | 522 | 8.12 | 141 | 506 | 538 |
| Cues + Nutrients | 525 | 8.12 | 141 | 509 | 541 |
| Cues | 531 | 8.12 | 141 | 515 | 547 |
| ASW | 530 | 8.12 | 141 | 513 | 546 |
| Contrasts | Estimate | SE | df | t.ratio | p.value |
| Nutrients – (cues + nutrients) | -3.016 | 5.44 | 297 | -0.555 | 0.9453 |
| Nutrients – Cues | -8.919 | 5.44 | 297 | -1.640 | 0.3579 |
| Nutrients – ASW | -7.826 | 5.44 | 297 | -1.439 | 0.4760 |
| (cues + nutrients) – Cues | -5.903 | 5.44 | 297 | -1.085 | 0.6988 |
| (cues + nutrients) – ASW | -4.810 | 5.44 | 297 | -0.884 | 0.8129 |
| Cues - ASW | 1.092 | 5.44 | 297 | 0.201 | 0.9971 |
| **Experiment Day = 12** | | | | | |
| Resuspended media | EMM | SE | df | Lower CL | Upper CL |
| Nutrients | 538 | 8.12 | 141 | 522 | 554 |
| Cues + Nutrients | 548 | 8.12 | 141 | 532 | 564 |
| Cues | 556 | 8.12 | 141 | 540 | 572 |
| ASW | 554 | 8.12 | 141 | 537 | 570 |
| Contrasts | Estimate | SE | df | t.ratio | p.value |
| Nutrients – (cues + nutrients) | -10.066 | 5.44 | 297 | -1.851 | 0.2518 |
| Nutrients – Cues | -18.253 | 5.44 | 297 | -3.356 | 0.0049 |
| Nutrients – ASW | -15.462 | 5.44 | 297 | -2.843 | 0.0245 |
| (cues + nutrients) – Cues | -8.187 | 5.44 | 297 | -1.505 | 0.4356 |
| (cues + nutrients) – ASW | -5.396 | 5.44 | 297 | -0.992 | 0.7540 |
| Cues - ASW | 2.790 | 5.44 | 297 | 0.513 | 0.9559 |
| **Experiment Day = 14** | | | | | |
| Resuspended media | EMM | SE | df | Lower CL | Upper CL |
| Nutrients | 543 | 8.12 | 141 | 526 | 559 |
| Cues + Nutrients | 553 | 8.12 | 141 | 537 | 569 |
| Cues | 554 | 8.12 | 141 | 538 | 570 |
| ASW | 553 | 8.12 | 141 | 537 | 569 |
| Contrasts | Estimate | SE | df | t.ratio | p.value |
| Nutrients – (cues + nutrients) | -10.624 | 5.44 | 297 | -1.953 | 0.2083 |
| Nutrients – Cues | -11.410 | 5.44 | 297 | -2.098 | 0.1561 |
| Nutrients – ASW | -10.823 | 5.44 | 297 | -1.990 | 0.1940 |
| (cues + nutrients) – Cues | -0.786 | 5.44 | 297 | -0.145 | 0.9989 |
| (cues + nutrients) – ASW | -0.200 | 5.44 | 297 | -0.037 | 1.0000 |
| Cues - ASW | 0.586 | 5.44 | 297 | 0.108 | 0.9996 |

**Table S12.** Linear mixed effect models and post-hoc comparisons testing the relationship between average cell volume ($\mu m^3$) and experiment day in the different media treatments (Resuspended media, 4 levels) for the algal species *Nannochloropsis granulata* (3 hours exposure). "Flask_ID" identifying the parent population was included as a random effect in the model. Exp Day = Experiment Day. Vol = Average cell volume. EMM = estimated marginal means. CL = 95% confidence level.

| ***Nannochloropsis* Average cell volume** | | | | | | |
|---|---|---|---|---|---|---|
| | Sum Sq | Mean Sq | NumDF | DenDF | F value | Pr (>F) |
| Exp Day | 119.23 | 10.839 | 11 | 108 | 13.301 | $1.084 \times 10^{-15}$ (***) |
| Resuspended media | 429.51 | 143.171 | 3 | 324 | 175.690 | $<2.2 \times 10^{-16}$ (***) |
| Exp Day × Resuspended media | 280.51 | 8.500 | 33 | 324 | 10.431 | $<2.2 \times 10^{-16}$ (***) |
| Post hoc test on experiment day × resuspended media | | | | | | |
| **Experiment Day = 1** | | | | | | |
| Resuspended media | EMM | SE | df | Lower CL | Upper CL | |
| Nutrients | 21 | 0.696 | 140 | 19.6 | 22.4 | |
| Cues + Nutrients | 22.5 | 0.696 | 140 | 21.1 | 23.8 | |
| Cues | 24 | 0.696 | 140 | 22.6 | 25.4 | |
| ASW | 25.6 | 0.696 | 140 | 24.2 | 27.0 | |
| Contrasts | Estimate | SE | df | t.ratio | p.value | |
| Nutrients – (cues + nutrients) | -1.4356 | 0.404 | 324 | -3.556 | 0.0024 | |
| Nutrients – Cues | -2.9755 | 0.404 | 324 | -7.37 | <0.0001 | |
| Nutrients – ASW | -4.5942 | 0.404 | 324 | -11.38 | <0.0001 | |
| (cues + nutrients) – Cues | -1.5399 | 0.404 | 324 | -3.814 | 0.0009 | |
| (cues + nutrients) – ASW | -3.1586 | 0.404 | 324 | -7.824 | <0.0001 | |
| Cues – ASW | -1.6187 | 0.404 | 324 | -4.009 | 0.0004 | |
| **Experiment Day = 3** | | | | | | |
| Resuspended media | EMM | SE | df | Lower CL | Upper CL | |
| Nutrients | 22.3 | 0.696 | 140 | 20.9 | 23.6 | |
| Cues + Nutrients | 23.6 | 0.696 | 140 | 22.2 | 24.9 | |
| Cues | 25.8 | 0.696 | 140 | 24.5 | 27.2 | |
| ASW | 23.9 | 0.696 | 140 | 22.6 | 25.3 | |
| Contrasts | Estimate | SE | df | t.ratio | p.value | |
| Nutrients – (cues + nutrients) | -1.2880 | 0.404 | 324 | -3.190 | 0.0084 | |
| Nutrients – Cues | -3.5678 | 0.404 | 324 | -8.837 | <0.0001 | |
| Nutrients – ASW | -1.6709 | 0.404 | 324 | -4.139 | 0.0003 | |
| (cues + nutrients) – Cues | -2.2798 | 0.404 | 324 | -5.647 | <0.0001 | |
| (cues + nutrients) – ASW | -0.3829 | 0.404 | 324 | -0.948 | 0.7787 | |
| Cues - ASW | 1.8969 | 0.404 | 324 | 4.699 | <0.0001 | |
| **Experiment Day = 4** | | | | | | |
| Resuspended media | EMM | SE | df | Lower CL | Upper CL | |
| Nutrients | 27.9 | 0.696 | 140 | 26.5 | 29.3 | |
| Cues + Nutrients | 27.9 | 0.696 | 140 | 26.6 | 29.3 | |
| Cues | 33.8 | 0.696 | 140 | 32.4 | 35.2 | |
| ASW | 29.6 | 0.696 | 140 | 28.2 | 31.0 | |
| Contrasts | Estimate | SE | df | t.ratio | p.value | |
| Nutrients – (cues + nutrients) | -0.0293 | 0.404 | 324 | -0.073 | 0.9999 | |
| Nutrients – Cues | -5.9046 | 0.404 | 324 | -14.626 | <0.0001 | |
| Nutrients – ASW | -1.6828 | 0.404 | 324 | -4.168 | 0.0002 | |
| (cues + nutrients) – Cues | -5.8752 | 0.404 | 324 | -14.553 | <0.0001 | |
| (cues + nutrients) – ASW | -1.6535 | 0.404 | 324 | -4.096 | 0.0003 | |
| Cues - ASW | 4.2218 | 0.404 | 324 | 10.457 | <0.0001 | |
| **Experiment Day = 5** | | | | | | |
| Resuspended media | EMM | SE | df | Lower CL | Upper CL | |
| Nutrients | 23.7 | 0.696 | 140 | 22.3 | 25.1 | |

| | | | | | |
|---|---|---|---|---|---|
| Cues + Nutrients | 26 | 0.696 | 140 | 24.6 | 27.4 |
| Cues | 28.4 | 0.696 | 140 | 27.0 | 29.8 |
| ASW | 25.4 | 0.696 | 140 | 24.1 | 26.8 |
| Contrasts | Estimate | SE | df | t.ratio | p.value |
| Nutrients – (cues + nutrients) | -2.2840 | 0.404 | 324 | -5.658 | <0.0001 |
| Nutrients – Cues | -4.6961 | 0.404 | 324 | -11.632 | <0.0001 |
| Nutrients – ASW | -1.7279 | 0.404 | 324 | -4.280 | 0.0001 |
| (cues + nutrients) – Cues | -2.4121 | 0.404 | 324 | -5.975 | <0.0001 |
| (cues + nutrients) – ASW | 0.5561 | 0.404 | 324 | 1.378 | 0.5144 |
| Cues - ASW | 2.9682 | 0.404 | 324 | 7.352 | <0.0001 |
| **Experiment Day = 7** | | | | | |
| Resuspended media | EMM | SE | df | Lower CL | Upper CL |
| Nutrients | 22.8 | 0.696 | 140 | 21.4 | 24.1 |
| Cues + Nutrients | 23.0 | 0.696 | 140 | 21.6 | 24.4 |
| Cues | 24.5 | 0.696 | 140 | 23.2 | 25.9 |
| ASW | 24.4 | 0.696 | 140 | 23.0 | 25.8 |
| Contrasts | Estimate | SE | df | t.ratio | p.value |
| Nutrients – (cues + nutrients) | -0.2369 | 0.404 | 324 | -0.587 | 0.9360 |
| Nutrients – Cues | -1.7606 | 0.404 | 324 | -4.361 | 0.0001 |
| Nutrients – ASW | -1.6224 | 0.404 | 324 | -4.019 | 0.0004 |
| (cues + nutrients) – Cues | -1.5237 | 0.404 | 324 | -3.774 | 0.0011 |
| (cues + nutrients) – ASW | -1.3855 | 0.404 | 324 | -3.432 | 0.0038 |
| Cues - ASW | 0.1382 | 0.404 | 324 | 0.342 | 0.9862 |
| **Experiment Day = 8** | | | | | |
| Resuspended media | EMM | SE | df | Lower CL | Upper CL |
| Nutrients | 21.6 | 0.696 | 140 | 20.2 | 23.0 |
| Cues + Nutrients | 21.3 | 0.696 | 140 | 19.9 | 22.7 |
| Cues | 23.3 | 0.696 | 140 | 21.9 | 24.7 |
| ASW | 23.9 | 0.696 | 140 | 22.5 | 25.3 |
| Contrasts | Estimate | SE | df | t.ratio | p.value |
| Nutrients – (cues + nutrients) | 0.2891 | 0.404 | 324 | 0.716 | 0.8906 |
| Nutrients – Cues | -1.7148 | 0.404 | 324 | -4.248 | 0.0002 |
| Nutrients – ASW | -2.3362 | 0.404 | 324 | -5.787 | <0.0001 |
| (cues + nutrients) – Cues | -2.0040 | 0.404 | 324 | -4.964 | <0.0001 |
| (cues + nutrients) – ASW | -2.6253 | 0.404 | 324 | -6.503 | <0.0001 |
| Cues - ASW | -0.6214 | 0.404 | 324 | -1.539 | 0.4153 |
| **Experiment Day = 9** | | | | | |
| Resuspended media | EMM | SE | df | Lower CL | Upper CL |
| Nutrients | 21.6 | 0.696 | 140 | 20.2 | 22.9 |
| Cues + Nutrients | 21 | 0.696 | 140 | 19.6 | 22.4 |
| Cues | 23 | 0.696 | 140 | 21.6 | 24.4 |
| ASW | 22.8 | 0.696 | 140 | 21.4 | 24.1 |
| Contrasts | Estimate | SE | df | t.ratio | p.value |
| Nutrients – (cues + nutrients) | 0.5759 | 0.404 | 324 | 1.426 | 0.4837 |
| Nutrients – Cues | -1.4312 | 0.404 | 324 | -3.545 | 0.0025 |
| Nutrients – ASW | -1.1965 | 0.404 | 324 | -2.964 | 0.0171 |
| (cues + nutrients) – Cues | -2.0071 | 0.404 | 324 | -4.972 | <0.0001 |
| (cues + nutrients) – ASW | -1.7724 | 0.404 | 324 | -4.390 | 0.0001 |
| Cues - ASW | 0.2346 | 0.404 | 324 | 0.581 | 0.9377 |
| **Experiment Day = 10** | | | | | |
| Resuspended media | EMM | SE | df | Lower CL | Upper CL |
| Nutrients | 22.2 | 0.696 | 140 | 20.8 | 23.6 |
| Cues + Nutrients | 22.2 | 0.696 | 140 | 20.9 | 23.6 |
| Cues | 24 | 0.696 | 140 | 22.6 | 25.4 |
| ASW | 23.7 | 0.696 | 140 | 22.3 | 25 |
| Contrasts | Estimate | SE | df | t.ratio | p.value |
| Nutrients – (cues + nutrients) | -0.0291 | 0.404 | 324 | -0.072 | 0.9999 |
| Nutrients – Cues | -1.8042 | 0.404 | 324 | -4.469 | 0.0001 |
| Nutrients – ASW | -1.4569 | 0.404 | 324 | -3.609 | 0.0020 |

| | | | | | |
|---|---|---|---|---|---|
| (cues + nutrients) – Cues | -1.7751 | 0.404 | 324 | -4.397 | 0.0001 |
| (cues + nutrients) – ASW | -1.4278 | 0.404 | 324 | -3.537 | 0.0026 |
| Cues - ASW | 0.3473 | 0.404 | 324 | 0.860 | 0.8253 |
| **Experiment Day = 11** | | | | | |
| Resuspended media | EMM | SE | df | Lower CL | Upper CL |
| Nutrients | 22.6 | 0.696 | 140 | 21.2 | 23.9 |
| Cues + Nutrients | 22.8 | 0.696 | 140 | 21.4 | 24.2 |
| Cues | 23.3 | 0.696 | 140 | 21.9 | 24.7 |
| ASW | 23.5 | 0.696 | 140 | 22.2 | 24.9 |
| Contrasts | Estimate | SE | df | t.ratio | p.value |
| Nutrients – (cues + nutrients) | -2375 | 0.404 | 324 | -0.588 | 0.9355 |
| Nutrients – Cues | -0.7438 | 0.404 | 324 | -1.842 | 0.2554 |
| Nutrients – ASW | -0.9742 | 0.404 | 324 | -2.413 | 0.0766 |
| (cues + nutrients) – Cues | -0.5063 | 0.404 | 324 | -1.254 | 0.5929 |
| (cues + nutrients) – ASW | -0.7367 | 0.404 | 324 | -1.825 | 0.2635 |
| Cues - ASW | -0.2304 | 0.404 | 324 | -0.571 | 0.9407 |
| **Experiment Day = 14** | | | | | |
| Resuspended media | EMM | SE | df | Lower CL | Upper CL |
| Nutrients | 22.9 | 0.696 | 140 | 21.5 | 24.2 |
| Cues + Nutrients | 22.3 | 0.696 | 140 | 20.9 | 23.6 |
| Cues | 23.8 | 0.696 | 140 | 22.4 | 25.2 |
| ASW | 24 | 0.696 | 140 | 22.6 | 25.4 |
| Contrasts | Estimate | SE | df | t.ratio | p.value |
| Nutrients – (cues + nutrients) | 0.6099 | 0.404 | 324 | 1.511 | 0.4322 |
| Nutrients – Cues | -0.9111 | 0.404 | 324 | -2.257 | 0.1105 |
| Nutrients – ASW | -1.1123 | 0.404 | 324 | -2.755 | 0.0314 |
| (cues + nutrients) – Cues | -1.5211 | 0.404 | 324 | -3.768 | 0.0011 |
| (cues + nutrients) – ASW | -1.7222 | 0.404 | 324 | -4.266 | 0.0002 |
| Cues - ASW | -0.2012 | 0.404 | 324 | -0.498 | 0.9594 |
| **Experiment Day = 16** | | | | | |
| Resuspended media | EMM | SE | df | Lower CL | Upper CL |
| Nutrients | 24 | 0.696 | 140 | 22.6 | 25.4 |
| Cues + Nutrients | 23.7 | 0.696 | 140 | 22.3 | 25.1 |
| Cues | 24.9 | 0.696 | 140 | 23.6 | 26.3 |
| ASW | 25.1 | 0.696 | 140 | 23.7 | 26.5 |
| Contrasts | Estimate | SE | df | t.ratio | p.value |
| Nutrients – (cues + nutrients) | 0.2779 | 0.404 | 324 | 0.688 | 0.9015 |
| Nutrients – Cues | -0.9425 | 0.404 | 324 | -2.335 | 0.0924 |
| Nutrients – ASW | -1.0875 | 0.404 | 324 | -2.694 | 0.0371 |
| (cues + nutrients) – Cues | -1.2205 | 0.404 | 324 | -3.023 | 0.0143 |
| (cues + nutrients) – ASW | -1.3655 | 0.404 | 324 | -3.382 | 0.0045 |
| Cues - ASW | -0.1450 | 0.404 | 324 | -0.359 | 0.9841 |
| **Experiment Day = 18** | | | | | |
| Resuspended media | EMM | SE | df | Lower CL | Upper CL |
| Nutrients | 27.2 | 0.696 | 140 | 25.9 | 28.6 |
| Cues + Nutrients | 28.3 | 0.696 | 140 | 26.9 | 29.6 |
| Cues | 28.4 | 0.696 | 140 | 27.1 | 29.8 |
| ASW | 28.8 | 0.696 | 140 | 27.5 | 30.2 |
| Contrasts | Estimate | SE | df | t.ratio | p.value |
| Nutrients – (cues + nutrients) | -1.0318 | 0.404 | 324 | -2.556 | 0.0536 |
| Nutrients – Cues | -1.2028 | 0.404 | 324 | -2.979 | 0.0163 |
| Nutrients – ASW | -1.6209 | 0.404 | 324 | -4.015 | 0.0004 |
| (cues + nutrients) – Cues | -0.1710 | 0.404 | 324 | -0.424 | 0.9744 |
| (cues + nutrients) – ASW | -0.5891 | 0.404 | 324 | -1.459 | 0.4634 |
| Cues - ASW | -0.4181 | 0.404 | 324 | -1.036 | 0.7286 |
